# PRIME: scalable, robust inference of mechanistic cell states from multimodal single-cell counts via probability generating functions

**DOI:** 10.64898/2026.06.04.730253

**Authors:** Shiyue Li, Yiling Wang, Qingchao Jiang, Ramon Grima, Edward Z. Cao

## Abstract

Single-cell multiomic technologies can now quantify complementary RNA species within the same cell, creating an opportunity to move beyond descriptive clustering toward mechanistically interpretable cell states. Yet most current methods depend on heuristic integration steps and become computationally burdensome at scale, limiting their ability to robustly detect subtle kinetic differences across heterogeneous populations. Here we introduce PRIME, a scalable framework for mechanistic cell-state discovery from multimodal single-cell count data. PRIME embeds multimodal measurements in a probability generating function (PGF) space, where transcriptional dynamics are encoded compactly and compared efficiently. This representation enables robust inference of latent kinetic structure and supports rapid cell grouping with a power K-means backbone that remains stable under noise, sparsity, and multimodality. Across synthetic benchmarks and experimental multimodal datasets, PRIME consistently recovers cell populations distinguished by transcriptional kinetics, outperforms conventional integration-and-clustering pipelines in robustness, and yields interpretable parameters that link observed variability to underlying regulatory mechanisms. By providing a mathematically principled yet practical route from multimodal counts to kinetic cell states, PRIME empowers biologists to uncover dynamic transcriptional regimes, dissect regulatory heterogeneity, and connect cell identity to mechanism rather than markers.

## 1. INTRODUCTION

Single-cell genomics has transformed modern biology by allowing researchers to measure gene activity in individual cells at an unprecedented scale and level of detail [1–4]. This has made it possible to identify many distinct cell states and types within complex tissues. However, most current analysis methods remain largely descriptive: cells are grouped according to similarities in their gene expression profiles, and these groups are then interpreted using marker genes and downstream statistical analyses [5–7]. While this approach is highly effective for organizing and annotating data, it does not directly explain the biological mechanisms that produce variability between cells or drive transitions from one state to another.

Multimodal single-cell technologies bring this challenge into sharper focus, while also offering a way forward. In particular, methods that measure multiple RNA species in the same cell — most notably unspliced and spliced transcripts, or nascent and mature RNA — provide access to transcriptional dynamics that is inaccessible to single-modality snapshots. These measurements have already transformed the study of cell-fate dynamics, most prominently through RNA velocity and its subsequent extensions, which use RNA processing states to infer short-timescale changes in cell state [8–15]. In parallel, metabolic labeling and time-resolved single-cell approaches — including NASC-seq, scSLAM-seq, sci-fate, scNT-seq, scGRO-seq and deep new-RNA sequencing — have greatly expanded the experimental toolkit for measuring nascent transcription at scale [16–21]. Together, these advances make it increasingly clear that cell identity is not just a static expression profile, but a dynamic process that can be observed experimentally. This motivates a shift from expression-defined clusters toward *kinetically grounded cell states*, in which cells are grouped by shared regulatory and transcriptional regimes rather than by superficial similarity in observed counts.

Despite major advances in integration algorithms, extracting mechanistic structure from multimodal counts remains a computational and statistical challenge. Widely used frameworks including weighted-nearest-neighbor strategies and deep generative models have substantially improved multimodal alignment, representation learning, and downstream clustering [6, 22–25]. Recent community-scale benchmarking further underscores both the breadth of integration approaches and the difficulty of selecting methods that remain robust across datasets, tasks, and modality combinations [7, 26, 27]. However, these methods are typically optimized for representation quality and cross-modality alignment, rather than for identifying cell states that are *explicitly defined by mechanistic parameters of transcriptional dynamics*. Consequently, mechanistic interpretability is usually recovered only after clustering, rather than being built directly into the inference objective.

A promising alternative is to define cell populations through generative biophysical models that directly connect multimodal observations to transcriptional kinetics. This direction was recently crystallized by mechanistic clustering approaches such as meK-means, which cluster cells in a space of inferred transcriptional parameters by leveraging the causal relationship between unspliced and spliced RNA counts [28]. More broadly, model-based frameworks illustrate the growing feasibility of mechanistic inference directly from single-cell sequencing data [29–34]. The mechanistic viewpoint is conceptually compelling: it turns clustering into the discovery of shared kinetic regimes, offering an interpretable definition of cell states grounded in regulatory dynamics rather than in purely geometric properties of an embedding. Yet, two obstacles limit broader adoption. First, mechanistic inference often becomes computationally demanding when scaled to modern datasets, especially when likelihood evaluation requires repeated numerical approximations or costly model fitting. Second, clustering in mechanistic parameter space can be fragile in practice: single-cell measurements are sparse, noisy, and variable in sequencing depth, and clustering objectives can be sensitive to initialization and local minima.

Here we present Probability-generating-function–based Robust Inference for Mechanistic Embedding (PRIME), a scalable framework for mechanism-first cell-state discovery built on two design principles. First, PRIME recasts multimodal single-cell count modeling in a probability generating function (PGF) space, where stochastic transcriptional dynamics can be represented compactly and compared efficiently [35–40]. This is not merely a mathematical convenience but a computational lever: PGFs provide a compact representation of stochastic transcription models with gene state switching, RNA production and splicing. They make it possible to compute and compare joint count distributions analytically, while retaining both the dependence between modalities and the variability generated by intrinsic and extrinsic sources of noise [35–39]. By operating in PGF space, PRIME constructs a mechanistically faithful representation that captures features often blurred in conventional latent embeddings.

Second, PRIME performs cell grouping with a power K-means backbone rather than classical K-means [41]. This choice is critical because standard K-means, while attractive for its simplicity and speed, is notoriously sensitive to initialization and poor local minima, and these weaknesses are amplified in high-dimensional, noisy single-cell settings. In PRIME, this additional optimization robustness is essential because the differences between underlying transcriptional kinetics are often small compared with the sparsity and depth variability of single-cell measurements. Stable optimization is therefore critical for identifying reproducible kinetic states rather than dataset-specific partitions. The result is a practical and interpretable framework for identifying cell states through transcriptional dynamics. Instead of combining multimodal measurements by heuristic weighting or alignment, PRIME treats them jointly as coupled observations generated by the same underlying stochastic process. This distinction matters biologically: two populations may exhibit similar mean expression yet be driven by fundamentally different kinetic programs, for example, differences in burst frequency (how often transcriptional bursts occur), burst size (how many transcripts are produced per burst), RNA processing kinetics, or effective degradation, whereas cells that appear far apart in expression space may nonetheless share a common underlying kinetic architecture. PRIME is designed to resolve such regimes by defining cell identity through kinetic structure, not markers alone.

## 2. RESULTS

### 2.1. Simple PGF transformation substantially improves the accuracy of identifying cell states

To motivate the development of PRIME and illustrate the advantages of computation in PGF space, we performed a comparative analysis on five single-cell RNA-sequencing datasets: a human skin dataset [42], a mouse lung dataset [42], a mouse primary motor cortex dataset [43], and two human lung adenocarcinoma datasets [44]. These datasets span multiple organisms, tissues, cell-type compositions, and sequencing technologies, thereby providing a broad testbed for representation robustness. Each dataset was processed into two count matrices: the unspliced matrix **U** = [*U*_*gi*_] and the spliced matrix **S** = [*S*_*gi*_], where *U*_*gi*_ and *S*_*gi*_ denote the numbers of unspliced and spliced transcripts of gene *g* in cell *i*, respectively. Highly variable genes (HVGs) were selected following the standard protocol in Ref. [28]. Details of data acquisition, preprocessing, and HVG selection are provided in Methods.

We compared two strategies for constructing cell-level feature representations. (i) Count-based representation. For each cell *i*, we concatenated the unspliced and spliced counts of all HVGs into a single vector *y*_*i*_. This corresponds to the conventional approach widely used in multimodal or multiomic integration. (ii) PGF-based representation. For a given gene *g*, the standard empirical probability generating function (EPGF) over a cell population is

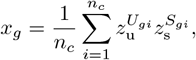

which summarizes the distribution of unspliced and spliced counts across *n*_*c*_ cells [45]. However, this population-level EPGF is not itself a cell-level feature representation. To retain cell-specific information, we instead associated each gene–cell pair (*g, i*) with the EPGF of the observed count pair {(*U*_*gi*_, *S*_*gi*_)}, viewed as a point mass at that observation

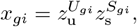

evaluated over a predefined grid with *z*_u_, *z*_s_ ∈ {0.5, 0.6, 0.7, 0.8, 0.9}. This produces a vector **x**_*gi*_ of length 25. Concatenating **x**_*gi*_ across all HVGs yields the PGF feature vector **x**_*i*_ for cell *i*. The construction of PGF-based cell representations is illustrated in Fig. 1a. Using both the conventional count-based features and the PGF-based features, we performed clustering with three algorithms that rely on distinct principles: K-means [46], Leiden [47], and fuzzy C-means [48] (implementation details in Supplementary Note 1). Because ground-truth annotations are available for all five datasets, clustering performance was quantified using the Adjusted Rand Index (ARI), a standard metric that measures agreement between inferred clusters and reference labels while correcting for chance.

**FIG. 1.**
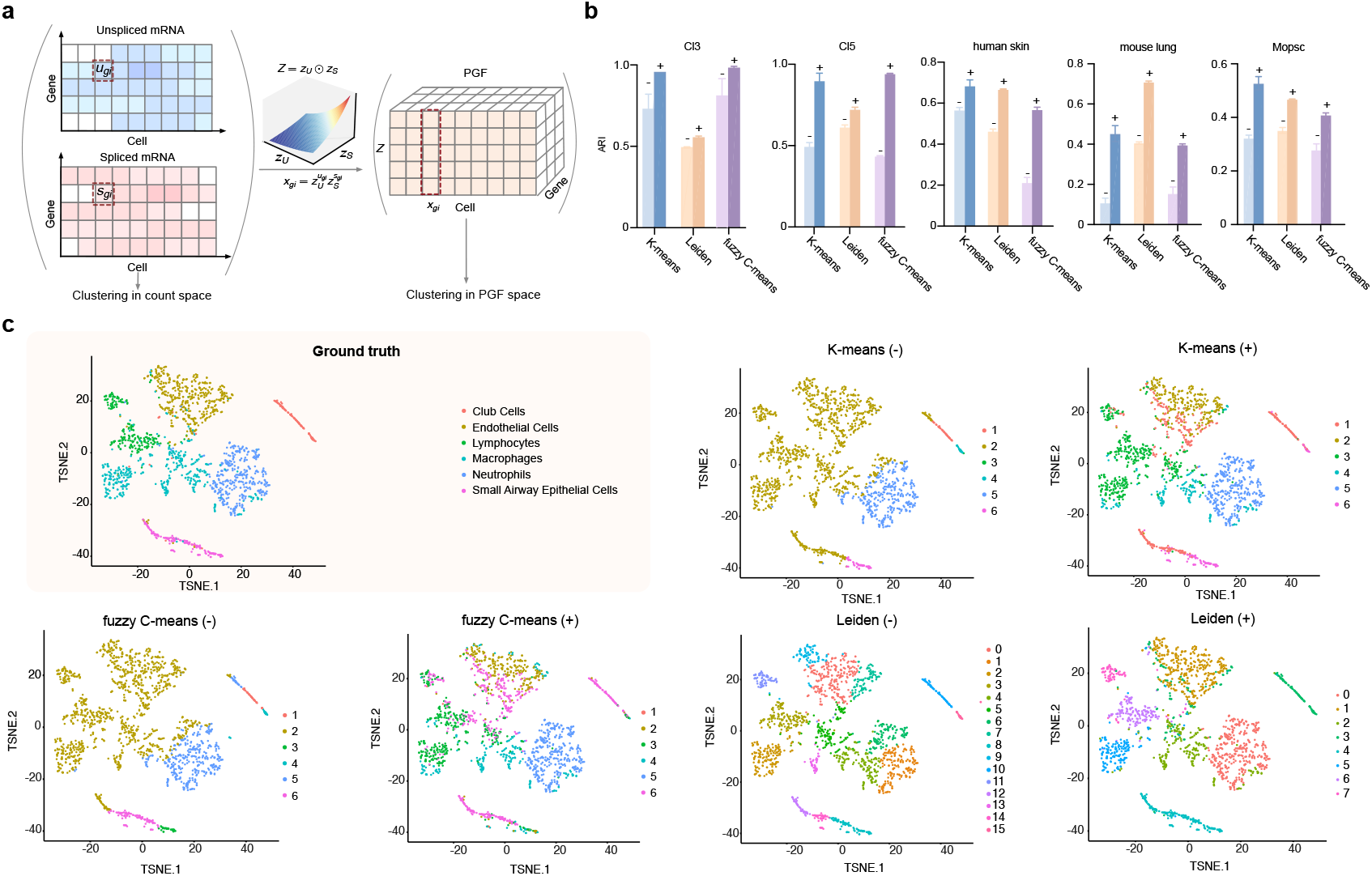
PGF transformation improves cell-state identification across datasets and clustering algorithms. (a) Schematic of feature construction. For each gene in each cell, paired unspliced and spliced counts are mapped to values of the empirical PGF on a predefined grid. Concatenation across highly variable genes yields a PGF-based representation of the cell. (b) Clustering accuracy measured by ARI for five datasets. Results are shown for three clustering algorithms (K-means, Leiden, fuzzy C-means) using conventional count-based features (“−”) or PGF-based features (“+”). For each method, the mean ARI and its standard error (SEM) were estimated from five independent runs; error bars indicate the SEM. PGF representations consistently achieve higher agreement with ground-truth annotations. (c) Two-dimensional visualization of the mouse lung dataset using t-SNE. PGF-based features yield clearer separation of major cell populations, including endothelial cells, lymphocytes, and macrophages.

Across datasets and algorithms, PGF-based representations (“+”) consistently achieved higher ARI scores than count-based representations (“−”), in some cases improving accuracy by up to twofold (Fig. 1b). Visualization of the mouse lung dataset using t-distributed stochastic neighbor embedding (t-SNE) further highlights clearer separation of major populations, including endothelial cells, lymphocytes, and macrophages, when PGF features are used (Fig. 1c). Gene expression is inherently stochastic [49, 50], so raw transcript counts provide a noisy and often unstable basis for defining cell state. Importantly, the motivation for PRIME is not that PGFs contain more information than an explicit probabilistic description; rather, they encode the same information in a more computationally useful form. Whereas explicit state-space formulations require manipulating large, sparse probability tables over joint count states, the PGF represents the same multimodal distribution as a compact functional object that can be evaluated and compared efficiently. This makes it possible to preserve not only mean abundance but also higher-order structure induced by bursting, processing and degradation, while providing a natural common representation for coupled RNA modalities. The benefit of PGF space is therefore not probabilistic content per se, but a smoother, more tractable representation for mechanistic inference and cell-state comparison.

### 2.2. Overview of the PRIME method

Building on the advantages of working in PGF space, we developed PRIME, a framework that learns transcriptional kinetics and cell states jointly from paired unspliced and spliced RNA counts. An overview of the workflow is shown in Fig. 2.

**FIG. 2.**
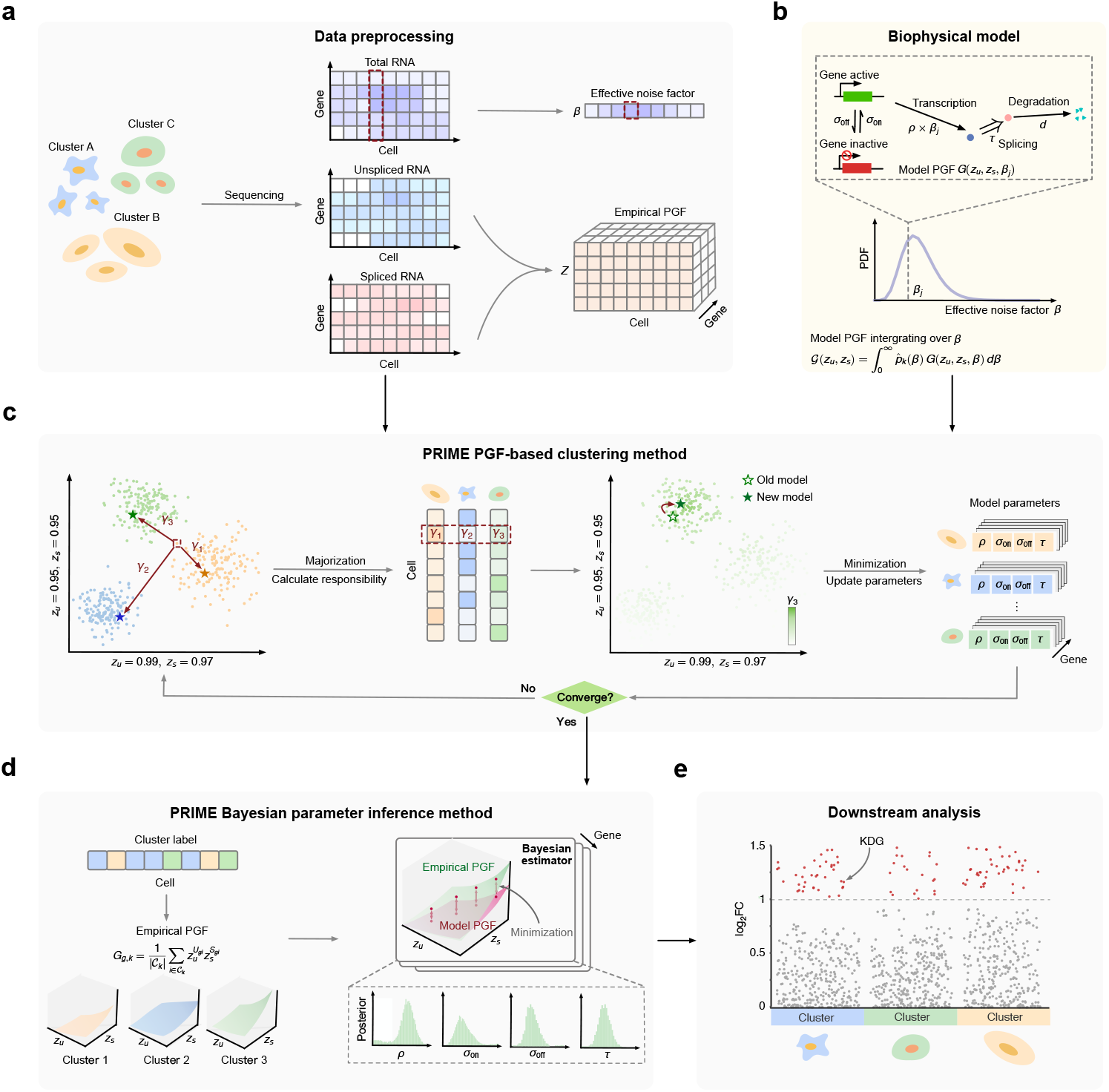
Overview of the PRIME framework. (a) Data preprocessing. From the unspliced and spliced RNA count matrices, empirical PGFs are computed for each gene in each cell. The effective noise factor *β* is estimated from total RNA counts to capture the combined effects of extrinsic variability (e.g., due to the coupling of transcription to cell volume) and technical noise. (b) Biophysical model. Transcription is described by a promoter-switching model in which the gene alternates between inactive and active states, producing unspliced transcripts that are converted into spliced molecules after a delay. This formulation admits an exact steady-state solution for the joint distribution of spliced and unspliced counts in PGF space. By integrating the analytical PGF solution over the distribution of *β*, the model accounts for intrinsic transcriptional noise, extrinsic variability, and sequencing noise. (c) PGF-based clustering. Using empirical PGFs of HVGs and the estimated *β* values, PRIME quantifies cell–cluster discrepancies in PGF space and performs soft clustering via a majorization–minimization algorithm that iteratively updates cluster responsibilities and cluster-specific kinetic parameters. (d) PGF-based Bayesian inference. Within each cluster, empirical PGFs are aggregated and used to infer transcriptional kinetic parameters using Bayesian inference, with the model PGF providing the likelihood. (e) Downstream analysis. PRIME enables discovery of cell states and identification of marker genes defined by changes in transcriptional kinetic parameters rather than differences in mean expression.

PRIME begins with a preprocessing step (Fig. 2a). For each gene in each cell, the observed unspliced and spliced counts are converted into an empirical PGF, following the construction in Fig. 1a. In parallel, we use Eq. (3) to estimate for each cell an effective noise factor, *β*, from the total RNA count matrix (the sum of unspliced and spliced counts across all genes). This quantity summarizes broad cell-to-cell variation due to the coupling of transcription to cell size [51–53] as well as technical effects such as variable capture efficiency and dropout [54–56]. Details of the estimation and physical interpretation of the effective noise factor are provided in Methods and Supplementary Note 2.

At the core of PRIME is a simple mechanistic description of transcription (Fig. 2b). In this model, a gene alternates between inactive and active states; when active, it produces unspliced RNA, which is subsequently processed into spliced RNA after a delay (Eq. (1)). A key feature of this formulation is that it can be expressed directly in PGF space (Eq. (2)), so model predictions can be evaluated in closed form rather than through repeated numerical solution of the full stochastic system. The model is further adapted to each cell through the effective noise factor *β*, allowing it to account for both biological variation and measurement noise. By integrating over the observed distribution of *β*, PRIME obtains a model representation that captures intrinsic transcriptional fluctuations, cell-to-cell variability and sequencing noise while remaining computationally efficient (Supplementary Note 2).

From this foundation, PRIME proceeds in two stages. In the first stage, cells are grouped in PGF space using highly variable genes together with their estimated noise factors (Fig. 2c). Rather than comparing cells directly in raw count space, PRIME casts cell clustering and kinetic-state learning as a joint optimization problem. In this formulation, the distance between a cell and a cluster is evaluated in PGF space (Eq. (4)), and these cell-to-cluster distances are combined through a power-mean objective (Eq. (5)). This objective generalizes standard clustering criteria and provides a smoother optimization landscape, improving robustness to noisy count-pair measurements and reducing sensitivity to poor local minima [41]. Optimization is performed with a majorization-minimization procedure, which extends the logic of soft-assignment clustering to the power-mean setting. In the majorization step, PRIME computes a responsibility for each cell-cluster pair, representing the degree of support for assigning a cell to a given cluster; these responsibilities are obtained directly from the gradient of the objective with respect to the corresponding cell-to-cluster distance (Eq. (6)). In the subsequent minimization step, cluster-specific kinetic parameters are updated in PGF space using these responsibilities as weights (Eq. (7)). Because this update is separable across genes and clusters, PRIME parallelizes the computation to maintain scalability on large datasets. The two steps are iterated until convergence, after which each cell is assigned to the cluster for which it has the highest responsibility. Cluster initialization is performed using standard K-means. Further technical details and derivations are provided in Methods and Supplementary Note 3.

In the second stage, PRIME performs gene-specific Bayesian inference within each cluster to quantify transcriptional kinetics and their uncertainty (Fig. 2d). This step builds on the fact that, when evaluated at a selected set of PGF coordinates, the empirical PGF can be treated as approximately multivariate Gaussian (Supplementary Note 4). This enables construction of a likelihood for the observed empirical PGF vector (Eq. (8)), whose mean is given by the model PGF solution (Eq. (2)) integrated over the estimated distribution of the effective noise factor *β* (Eqs. (9) and (11)), and whose covariance is computed directly from the data (Eq. (13)). With priors placed on the kinetic parameters, posterior inference yields cluster-specific estimates of transcriptional behaviour and their uncertainty; equivalently, it amounts to finding parameter values that minimize the covariance-weighted discrepancy between the observed empirical PGF and its model prediction (Fig. 2d). In this way, each cluster becomes an interpretable kinetic state, defined not only by its member cells but by the transcriptional program that best explains their joint unspliced and spliced RNA measurements. See Methods for more details.

These inferred kinetic states enable downstream analyses (Fig. 2e), including the identification of kinetically differentiated genes (KDGs), whose distinguishing features arise from changes in transcriptional dynamics rather than from differences in average expression alone. In this way, PRIME provides a mechanism-first route from multimodal count data to cell-state discovery, combining interpretable modeling, robust clustering and scalable inference within a single framework.

### 2.3. Benchmarking PRIME on synthetic datasets

To evaluate clustering accuracy, parameter recovery, and computational efficiency, we generated controlled synthetic datasets in which the ground-truth structure is known. The simulation workflow is summarized in Fig. S1. Briefly, each cell was assigned an effective noise factor *β* sampled from a Gamma distribution to mimic extrinsic and technical noise. We then specified the number of clusters, the number of HVGs, and the cluster sizes, after which gene-specific kinetic parameters of the transcription model (Eq. (1)) were drawn. Finally, unspliced and spliced transcript counts were generated in steady-state conditions using stochastic simulations implemented in `DelaySSAToolkit.jl` [57]. Full details of the protocol are provided in Supplementary Note 1.

Because the quadrature resolution (*n*_*z*_) and integration domain [*z*_min_, *z*_max_] are hyperparameters of PRIME, we performed a sensitivity analysis using a synthetic dataset with ten clusters. As shown in Fig. S2a, we evaluated *n*_*z*_ ∈ {5, 7, 9} and observed that *n*_*z*_ = 7 achieves stable performance without unnecessary computational overhead. We further examined integration ranges (0, 1), (0.5, 1), (0.7, 1), and (0.95, 1) (Fig. S2b). Restricting the domain to (0.95, 1) consistently yielded the highest clustering accuracy, indicating that the near-unity region of PGF space provides the most discriminative information. Unless otherwise specified, we therefore use *n*_*z*_ = 7 and the integration range (0.95, 1) throughout.

PRIME integrates several key design elements: representation of data in PGF space rather than raw count space, explicit correction of cell-to-cell variability through integration over the effective noise factor *β*, soft cluster assignments within a majorization-minimization optimization framework, and mechanistic guidance from a biophysical transcription model combined with a power K-means backbone. To assess the contribution of these components, we conducted an ablation study in which individual elements were selectively removed (Supplementary Note 1). This produced eight reduced variants of PRIME, which were compared with the full method on a synthetic dataset containing ten clusters. Clustering performance, summarized by the mean ARI and its SEM across five independent runs, is shown in Fig. 3a. The four reduced variants that operate directly in count space (Reduce 5-8) consistently underperform those based on PGF representations (Reduce 1-4), confirming that transforming the data into PGF space substantially improves the identifiability of cell states. Comparisons between PRIME and its hard-assignment counterparts (Reduce 1 and Reduce 2), which omit the power K-means backbone that naturally enables soft assignments, further highlight the importance of maintaining soft cluster responsibilities and explicitly modeling cell-to-cell variability. Removing either component leads to systematic reductions in ARI, indicating that probabilistic assignments and integration over extrinsic and technical noise stabilize the clustering landscape. Finally, comparison with the purely geometric approach in Reduce 3, which combines the PGF representation with the power K-means backbone but lacks mechanistic guidance, demonstrates the additional benefit of incorporating the biophysical transcription model. Including this mechanistic component yields further improvements in clustering accuracy beyond those achieved by PGF transformation alone.

**FIG. 3.**
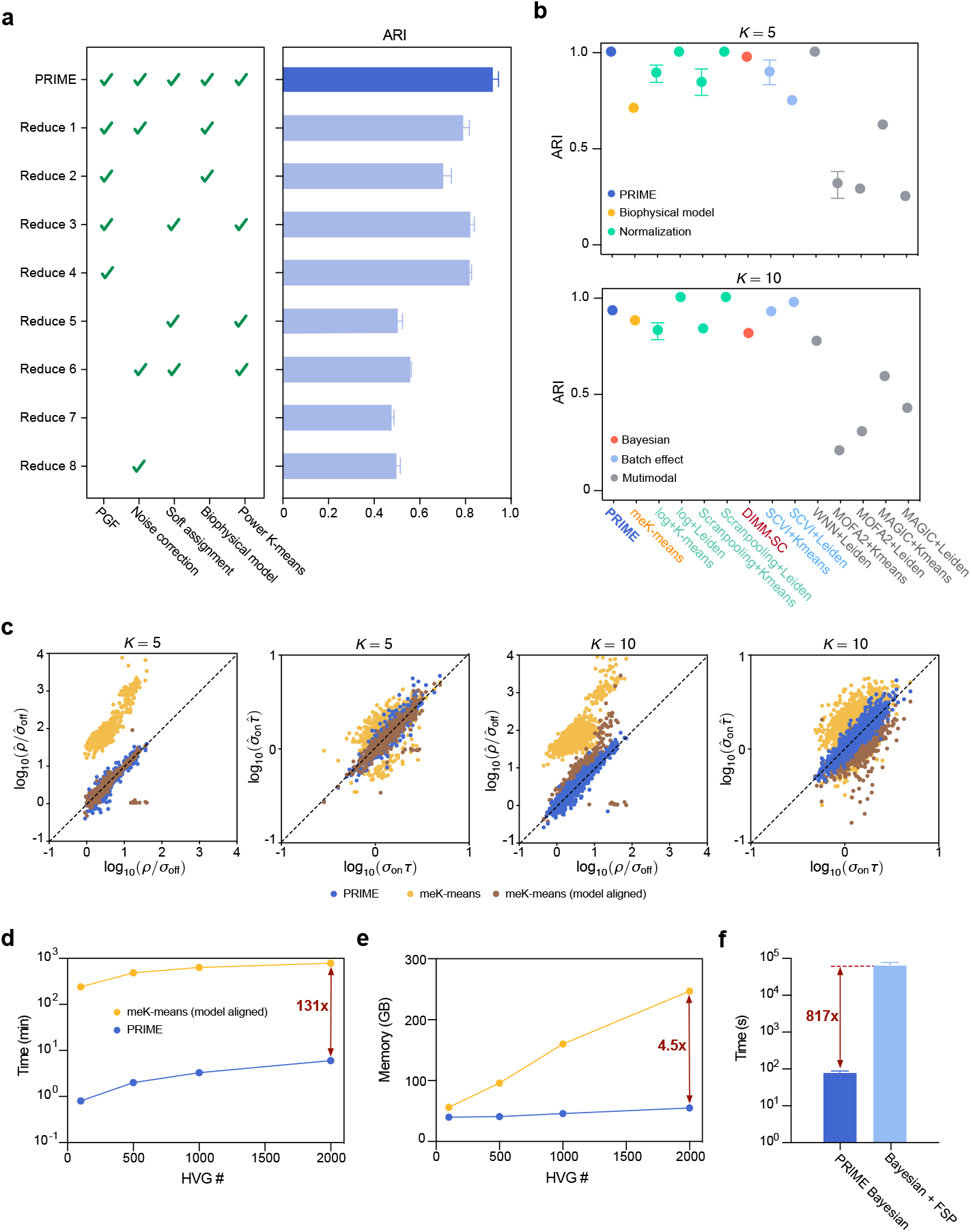
Benchmarking PRIME on synthetic datasets. (a) Ablation study on a 10-cluster synthetic dataset. PRIME is compared with eight reduced variants in which key components (mechanistic guidance, PGF representation, noise correction, soft assignment or power K-means backbone) are selectively removed. Bars represent the mean ARI across five independent runs; error bars denote the SEM. (b) Comparison of PRIME with representative clustering pipelines based on distinct methodological principles, including mechanistic modeling (meK-means), normalization-based pipelines, batch-correction approaches, multimodal integration methods, and Bayesian clustering frameworks. PRIME consistently ranks among the top-performing methods for datasets with *K* = 5 and *K* = 10 clusters. Dots represent the mean ARI across five independent runs; error bars denote the SEM. (c) Parameter inference accuracy for clusters of 1600 cells (*K* = 5) and 800 cells (*K* = 10). Ground-truth and inferred parameters are mapped to burst size (*ρ/σ*_off_) and burst frequency (*σ*_on_*τ*). PRIME accurately recovers both quantities across clusters, whereas meK-means without model alignment systematically overestimates burst size and exhibits greater variance in burst-frequency estimates. (d,e) Computational efficiency of the clustering module under matched implementation conditions. Runtime (d) and memory usage (e) are shown as functions of the number of HVGs. PRIME demonstrates substantially better scalability, achieving a ~ 131 *×* speedup and lower memory consumption than meK-means for 2000 HVGs. Dots represent the mean of three independent runs; the SEM is negligible. (f) Computational efficiency of Bayesian parameter inference. PRIME’s PGF-based inference is compared with a count-based Bayesian estimator in which likelihoods are computed using FSP. PRIME achieves up to a ~ 817 *×* speedup while maintaining accurate parameter recovery. Bars represent the mean computational time across 10 genes; error bars denote the SEM.

In Fig. 3b, we benchmark PRIME against several groups of state-of-the-art clustering methods that represent distinct methodological principles. First, because PRIME is mechanistically guided, we compare it with the only previously reported biophysical model-driven clustering framework, meK-means [28]. Second, PRIME explicitly accounts for confounded extrinsic and technical variability through integration over the effective noise factor *β*, which serves a normalization role. We therefore compare it with standard normalization-based pipelines, which mainly account for sequencing-related technical variability by converting discrete raw counts into continuous representations, including log-transformation and Scran pooling [58], each followed by K-means or Leiden clustering. Third, because the effective noise factor conceptually relates to batch correction, we also evaluate variants of K-means and Leiden integrated with SCVI [59]. Fourth, as PRIME naturally integrates multiomic information through its mechanistic PGF formulation, we compare it with representative multimodal integration methods (MAGIC [60], MOFA2 [61], and WNN [22]) combined with standard clustering algorithms. Finally, we include Bayesian clustering approaches such as DIMM-SC [62]. Implementation details are provided in Supplementary Note 1, and Fig. S3 summarizes the functional features of the compared methods. Comparisons were performed on both the previously generated 10-cluster dataset and an additional 5-cluster dataset to assess scalability with respect to the number of clusters. Across both settings, PRIME consistently ranks among the top-performing methods in clustering accuracy (Fig. 3b). Notably, PRIME outperforms meK-means in both cases, the latter being the most directly comparable framework due to its mechanistic foundation.

Next, we benchmark the parameter inference accuracy of PRIME against its most closely related competitor, meK-means, in Fig. 3c. To enable a fair comparison of inferred kinetic parameters, we map both the ground-truth parameters and PRIME’s posterior estimates to two dimensionless quantities:

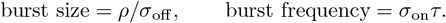

These quantities correspond, respectively, to the average number of mRNA molecules synthesized during each active transcriptional episode and to the activation frequency normalized by the splicing rate, and are directly estimated in meK-means. As shown in Fig. 3c, PRIME accurately recovers both burst size and burst frequency across clusters in both synthetic datasets. In contrast, meK-means systematically overestimates burst size and exhibits greater variance in burst-frequency estimates. This discrepancy mirrors the gap observed in clustering accuracy in Fig. 3b. Because the synthetic data were generated using the mechanistic model adopted by PRIME, we repeated the comparison after aligning the meK-means model with the data-generating model (see Supplementary Note 1 for details of the modifications to meK-means). Under this matched setting, the performance of meK-means improves. For *K* = 5, PRIME and meK-means achieve comparable parameter recovery and ARI (Fig. S4a). For *K* = 10, meK-means still underperforms PRIME in clustering accuracy (Fig. S4b), accompanied by a modest overestimation of burst size and underestimation of burst frequency. Together, these results highlight two key factors underlying PRIME’s performance: accurate mechanistic specification and parameter inference in PGF space.

To further assess computational efficiency under matched conditions, we compared PRIME’s clustering module with meK-means using an identical implementation environment, including the same programming language, optimization algorithm, and numerical tolerances (Supplementary Note 1). As shown in Fig. 3d and 3e, under these controlled settings the computational time and memory usage of both methods increase with the number of HVGs. However, the scaling behavior differs substantially. For 2000 HVGs, PRIME completes clustering in approximately 6 minutes, whereas meK-means requires nearly 13 hours, corresponding to a speedup of about 131×. Although PRIME operates in PGF space, which has higher dimensionality than the raw count space used by meK-means because each observed count pair (*U*_*gi*_, *S*_*gi*_) is represented by an empirical PGF evaluated at multiple (*z*_u_, *z*_s_) points, it nonetheless exhibits substantially lower memory usage. For 2000 HVGs, meK-means requires approximately 4.5 × more memory than PRIME. This difference arises because meK-means computes joint probabilities of model Eq. (1) using the Finite State Projection (FSP) method [63], which requires relatively large truncation sizes to maintain numerical accuracy and therefore increases memory consumption. These results demonstrate that representing single-cell data in PGF space improves not only clustering accuracy but also computational efficiency. This conclusion is further supported by a comparison between PRIME’s Bayesian parameter inference module and a Bayesian estimator based on count data whose likelihood is computed using FSP (Supplementary Note 1). As shown in Fig. 3f, the speedup is up to 817×, consistent with our previous findings in the context of transcriptional parameter inference without clustering [45].

### 2.4. Benchmarking PRIME on experimental datasets

We next applied PRIME to the five experimental datasets used in Fig. 1a and compared its clustering performance with 13 established methods. Because ground-truth kinetic parameters are not available for experimental data, evaluation was restricted to clustering accuracy. Among the five datasets, `human skin` and `mouse lung` exhibit explicit batch structure: the former aggregates samples from three healthy donors, and the latter from four mice under two experimental conditions. To ensure a fair comparison, we evaluated batch-aware pipelines when applicable. Specifically, ComBat [64] and MNN [65] were applied only when explicit batch annotations were available (Supplementary Note 1). In addition, SCVI [59] was included as a model-based batch correction approach that can account for batch effects without requiring predefined batch labels. As shown in Figs. 4a and S5, PRIME consistently ranks among the top-performing methods according to both ARI and normalized mutual information (NMI). These results suggest that the PGF-based mechanistic framework underlying PRIME naturally accounts for sources of variability such as extrinsic and technical noise, batch effects, and multimodal heterogeneity.

**FIG. 4.**
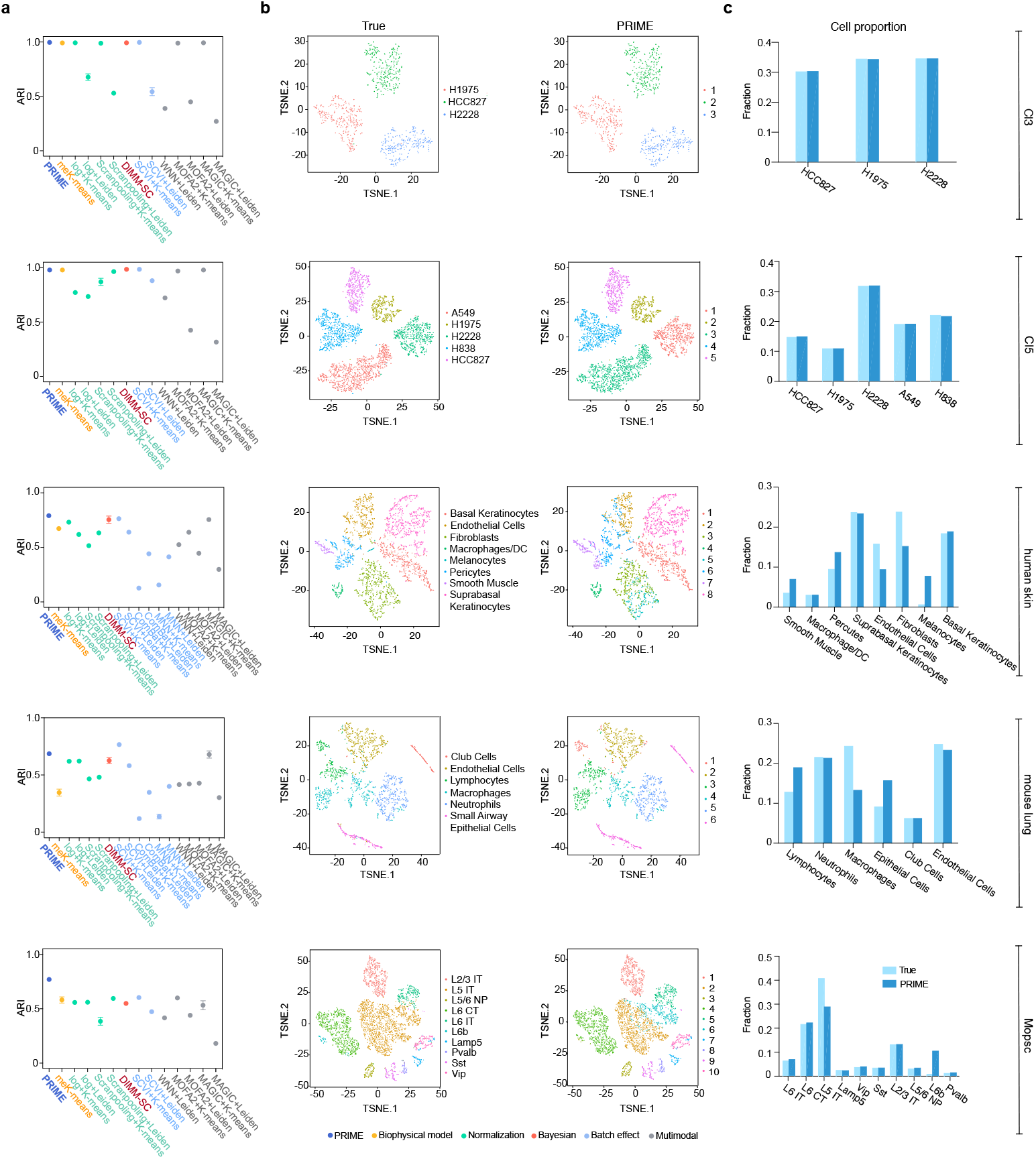
Performance of PRIME on experimental single-cell RNA-seq datasets. (a) Clustering accuracy across five experimental datasets. PRIME is compared with 13 state-of-the-art methods, including normalization-based (log transformation, Scran pooling), batch-corrected (ComBat, MNN, SCVI), multimodal-integration (MAGIC, MOFA2, WNN), Bayesian (DIMM-SC), and mechanistic (meK-means) approaches. Performance is quantified using ARI. Bars represent mean values across five independent runs; error bars denote SEM. (b) Two-dimensional visualization of clustering results in t-SNE space. Points are colored by cluster assignments inferred by PRIME and compared with ground-truth annotations for representative datasets. PRIME yields clear separation of major cell populations. (c) Breakdown of inferred clusters by annotated cell types. Rare populations (for example, L6 IT and Lamp5 neurons in `Mopsc`) are successfully resolved despite their low abundance. Together, these results demonstrate robust clustering performance of PRIME across heterogeneous biological systems and sequencing conditions.

A particularly informative comparison is with meK-means, the only previously reported biophysical model guided clustering framework. PRIME substantially outperforms meK-means on the `mouse lung` and `Mopsc` datasets. This difference is consistent with the distinct mechanistic assumptions underlying the two models. meK-means relies on a formulation in which negative binomial-like transcript distributions are attributed solely to transcriptional bursting, characterized by long inactive periods interspersed with brief high-activity episodes. However, recent theoretical work [66] demonstrates that negative binomial-like behavior can arise across broader regions of kinetic parameter space and is not uniquely diagnostic of canonical bursting. Constraining inference to a bursting-dominated regime can therefore bias estimates of burst size and burst frequency. Such parameter misspecification may propagate to clustering errors, particularly when a subset of HVGs does not conform to strongly bursty dynamics. Although more complex promoter models with multiple gene states have been proposed [67–69], it has been shown that, in the absence of time-resolved measurements, a two-state (telegraph-like) model is the minimal statistically identifiable description of transcription dynamics [45]. The telegraph-like model employed in PRIME spans the parameter regime assumed in meK-means while retaining analytical tractability in PGF space. This broader yet identifiable formulation likely contributes to the improved clustering robustness observed in experimental data.

Figure 4b visualizes PRIME’s clustering results relative to ground-truth annotations in t-SNE space. Using maximal alignment between inferred clusters and reference annotations (Supplementary Note 1), we quantified the composition of each inferred cluster and summarized the results in Fig. 4c. Overall, the inferred clusters show strong correspondence with known cell identities, indicating that PRIME preserves the major biological structure of the datasets. In the `mouse lung` dataset, cluster identities are largely consistent with the reference labels. The remaining discrepancies are localized; for example, a small fraction of macrophages is grouped with lymphocytes. Such misassignments likely reflect partial transcriptional similarity between immune cell states rather than a global clustering failure. A similar pattern is observed in the `Mopsc` dataset, where the L5 IT and L6b populations remain partially overlapping. This behavior is expected given the known transcriptomic continuum between these neuronal subtypes and the relatively small sample size of the L6b population, which limits statistical separability. PRIME accurately identifies even extremely rare populations in `Mopsc` with proportions below 0.05%. This demonstrates robustness to class imbalance and sensitivity to low-frequency cell states.

### 2.5. PRIME identifies kinetically differentiated genes beyond mean-expression analysis in mouse breast cancer

We first applied the PGF-based clustering module of PRIME to the mouse breast cancer dataset (`Brca`) reported in Ref. [70] and previously analyzed in Ref. [28]. Following the same experimental setting as in Ref. [28], we set the number of clusters to *K* = 5. The resulting clustering structure is visualized in Fig. 5a. To assess the biological consistency of the inferred clusters, we examined canonical marker genes previously reported to distinguish cell populations in this dataset. Figure 5b shows the mean expression patterns across clusters, quantified as the min–max normalized total transcript abundance (defined as the sum of unspliced and spliced counts and denoted as the scaled *µ*_total_). These values exhibit clear separation across the five clusters, confirming that PRIME recovers biologically meaningful cell states consistent with prior analyses.

**FIG. 5.**
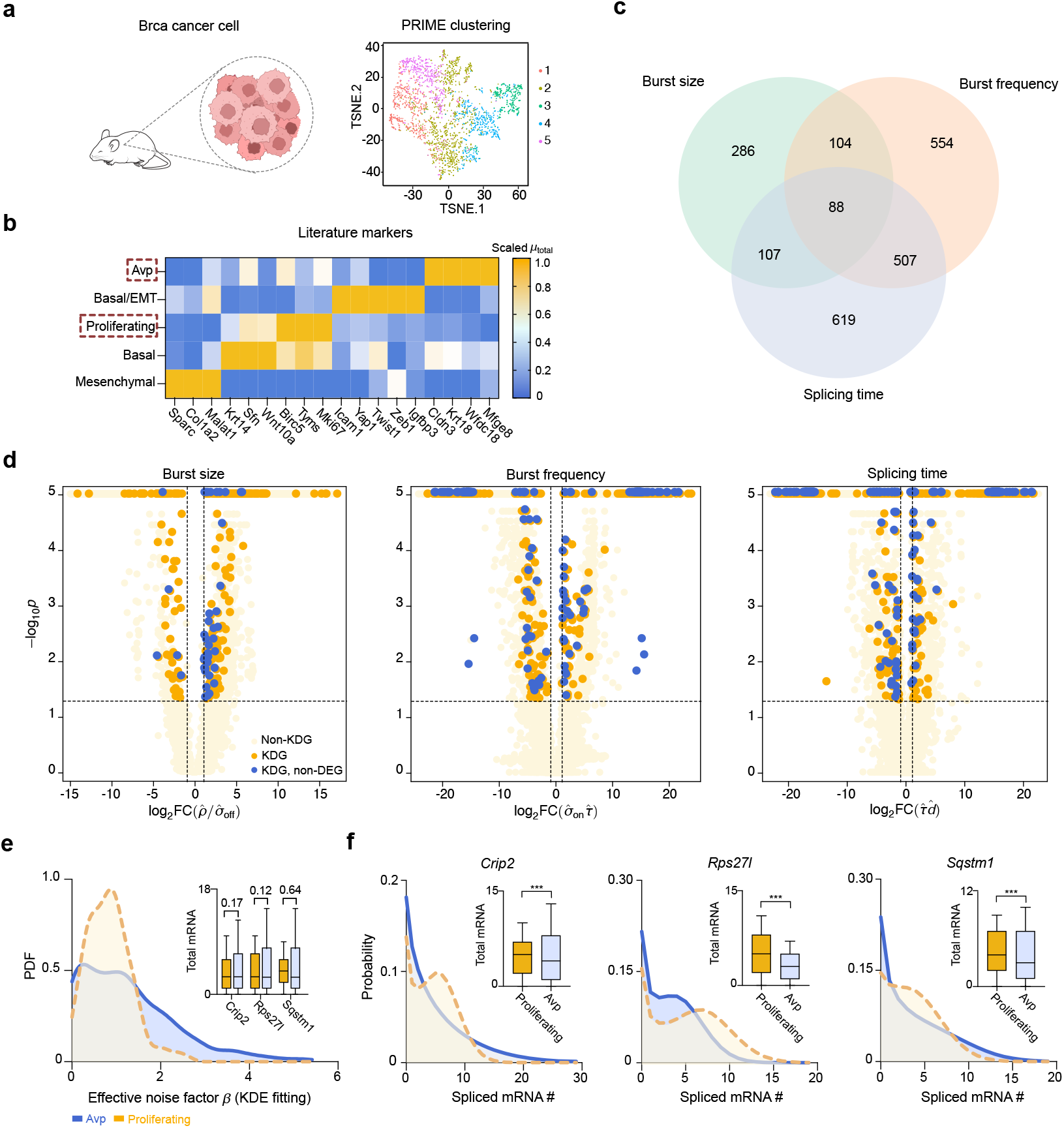
PRIME identifies kinetically differentiated genes beyond mean-expression analysis in mouse breast cancer data. (a) Visualization of PRIME clustering results for the mouse breast cancer (`Brca`) dataset with *K* = 5 clusters. Cells are colored by inferred cluster identity. (b) Heatmap of normalized mean expression (min-max normalized total RNA counts) for representative marker genes across the five clusters, confirming biologically coherent cell-state separation. (c) Summary of KDGs between two representative clusters (Avp and Proliferating) identified by PRIME based on inferred burst size, burst frequency, and splicing time. The Venn diagram illustrates the overlap and specificity among these three kinetic categories. (d) Volcano plots comparing the Avp and Proliferating clusters for the three kinetic parameters. The horizontal axis shows the log_2_ fold change of posterior median estimates, and the vertical axis shows the *p*-value associated with the fold change. Orange points indicate KDGs, whereas light yellow points denote non-KDGs. Blue points highlight KDGs that are not detected as DEGs based on mean expression. (e) Kernel density estimates of the inferred effective noise factor *β* for the Avp and Proliferating clusters, highlighting differences in extrinsic and technical variability between the two cell populations. Inset: mean expression levels of representative genes, showing limited separation when assessed using raw count-based averages. Labeled numbers indicate the corresponding *p*-values. (f) Model-reconstructed distributions of spliced RNA for representative KDGs (*Crip2, Rps27l*, and *Sqstm1*) after correcting for noise-factor effects by fixing *β* = 1. Insets show the corresponding reconstructed mean levels. The *p*-values in the insets were computed using two-sided Mann-Whitney tests applied to two sets of 1000 samples drawn from the reconstructed distributions using the inferred point estimates of kinetic parameters. Boxplots in the insets of (e,f) indicate the 10%, 25%, 50%, 75%, and 90% quantiles. ***: *p <* 10^−3^.

With reliable clustering established, we focused on the Avp and Proliferating cell populations (highlighted in Fig. 5b) and applied PRIME’s PGF-based Bayesian parameter inference framework together with the kinetic-differentiation analysis described in Methods. For each gene and cluster, posterior distributions of the kinetic parameters were inferred and subsequently transformed into three interpretable dimensionless quantities: burst size, burst frequency, and splicing time. Applying the KDG identification criteria across the whole genome yielded a substantial number of kinetically differentiated genes for each kinetic descriptor (Fig. 5c). The overlap and specificity among the three KDG categories are summarized by the Venn diagram in Fig. 5c, highlighting that different kinetic dimensions capture partially distinct aspects of transcriptional regulation.

Figure 5d presents volcano plots for burst size, burst frequency, and splicing time. KDGs that distinguish the Avp and Proliferating cell populations are highlighted in blue and orange, respectively. Notably, a subset of KDGs (blue) does not satisfy conventional differential-expression criteria based on mean transcript abundance (differentially expressed genes, DEGs; see Methods). This finding demonstrates that kinetic differentiation reveals regulatory differences that remain undetected by standard mean-expression analysis. More broadly, these results illustrate how integrating a mechanistic model with PGF-based inference enables a deeper characterization of cell-state differences beyond transcript abundance alone.

We further examined three representative KDGs *Crip2* (burst size), *Rps27l* (burst frequency), and *Sqstm1* (splicing time) to illustrate the mechanistic interpretation of the inferred parameters. Figure 5e shows the kernel-density-estimated distributions of the effective noise factor *β* for the Avp and Proliferating clusters, revealing clear differences in both shape and central tendency. The inset of Fig. 5e demonstrates that these genes exhibit minimal differences in mean expression, explaining why they are not detected as DEGs. However, when the inferred kinetic parameters are used to reconstruct the underlying transcript distributions (with *β* fixed to unity to remove extrinsic scaling effects), clear differences emerge in the predicted spliced-RNA distributions and their reconstructed means (Fig. 5f). This analysis highlights the importance of explicitly accounting for extrinsic and technical variability through the effective noise factor. Without such correction, substantial kinetic differences between cell populations may remain hidden, potentially obscuring biologically meaningful regulatory mechanisms.

Together, these results demonstrate that PRIME enables the identification of kinetically differentiated genes that cannot be detected by conventional expression-based analysis, providing a complementary and mechanistically grounded perspective on cellular heterogeneity.

### 2.6. PRIME identifies novel candidate marker genes for cutaneous T-cell lymphoma

We next applied PRIME to expanded CRISPR-compatible cellular indexing of transcriptomes and epitopes by sequencing (ECCITE-seq) data [71], which enable simultaneous measurement of transcript abundance and cell-surface protein levels. We focused on the `CTCL` and `ctrl` datasets, which contain peripheral blood mononuclear cells (PBMCs) collected from patients with cutaneous T-cell lymphoma and healthy donors, respectively (Methods).

To ensure robustness of downstream analyses, we first performed standard cell-level quality control. Cells were retained if the number of detected genes (defined as the number of non-zero entries in total RNA counts) was between 300 and 2500 and the fraction of mitochondrial transcripts was below 10%. HVGs were then selected following the protocol of Ref. [28] (Methods). The two datasets were processed independently but using the same preprocessing pipeline. For each dataset, effective noise factors were estimated according to Eq. (3), and PRIME PGF-based clustering was performed with the number of clusters fixed to *K* = 3, consistent with prior analysis [28]. Cluster identities were subsequently annotated using cell-surface protein markers (Supplementary Note 1). The resulting compositions of CD4^+^ T cells, CD8^+^ T cells, and monocytes across clusters are shown in Fig. 6a.

**FIG. 6.**
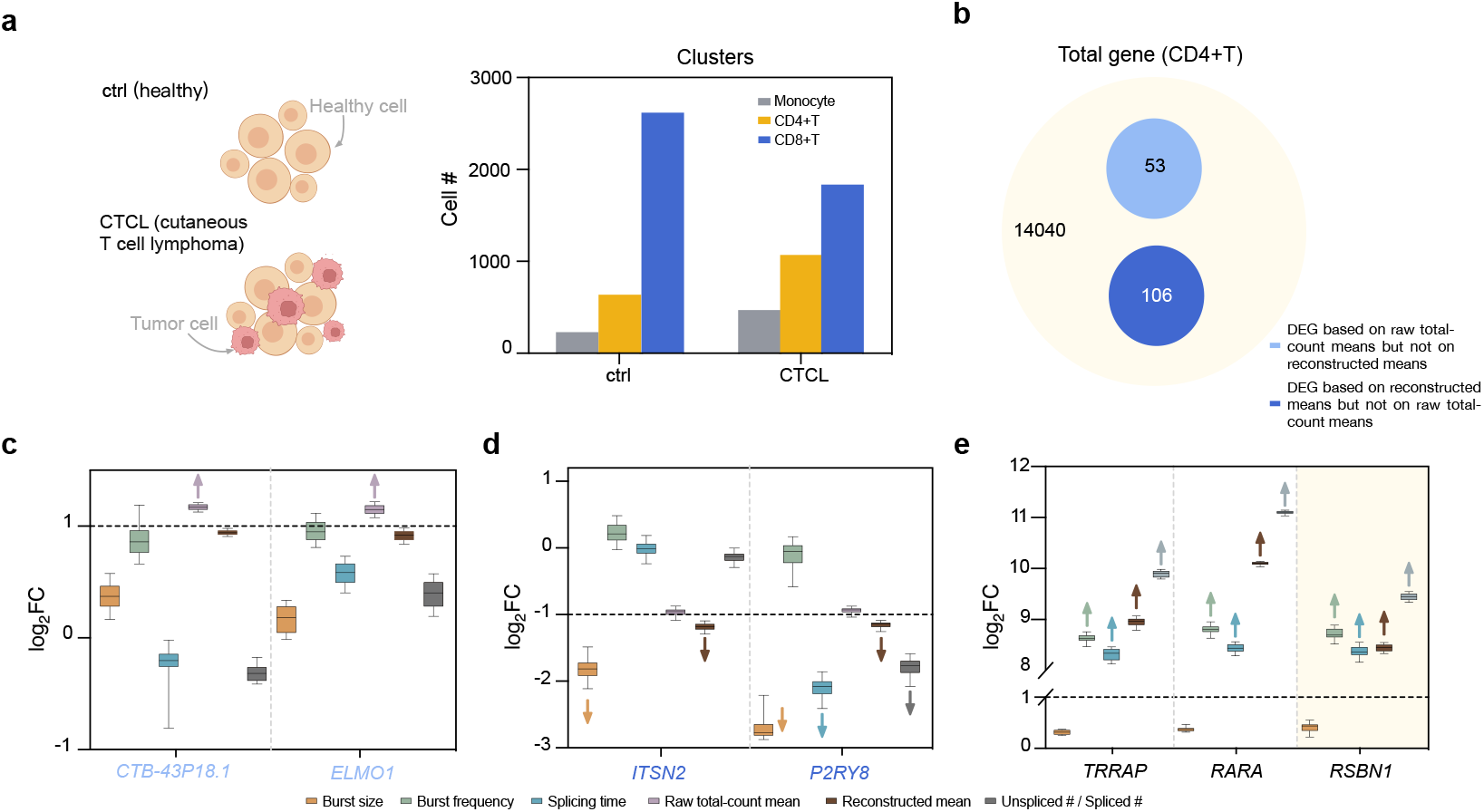
PRIME reveals novel marker genes in CTCL. (a) PRIME clustering of ECCITE-seq PBMC data from CTCL patients and healthy donors (ctrl). Cluster identities were assigned using cell-surface protein markers, revealing populations of CD4^+^ T cells, CD8^+^ T cells, and monocytes. (b) Comparison of differential-expression analyses based on raw total-count means and reconstructed means obtained from inferred kinetic parameters after correcting for the effective noise factor. The diagram summarizes genes uniquely detected by each approach. (c) Example genes with large fold changes in raw total-count means (|log_2_ FC| *>* 1) that become non-significant after reconstruction, consistent with unchanged kinetic parameters. (e) Example genes with modest raw-count differences but clear separation in reconstructed means, showing how kinetic inference reveals regulatory differences that are obscured in raw counts. (f) Kinetic signatures of representative CTCL marker genes. Reconstructed expression and inferred kinetic parameters for *TRRAP, RARA*, and the candidate marker *RSBN1* in the `CTCL` and `ctrl` samples. All three genes display a consistent regulatory pattern characterized by increased burst frequency, prolonged splicing time, elevated total RNA abundance, and shifts in the unspliced-to-spliced ratio. Boxplots in (c-e) indicate the 10%, 25%, 50%, 75%, and 90% quantiles. Arrows in (c–e) highlight significant changes in expression or kinetic parameters.

We then focused on CD4^+^ T cells and applied PRIME’s PGF-based Bayesian parameter inference framework to compare transcriptional kinetics between the `CTCL` and `ctrl` samples. A central aim of this analysis was to evaluate the impact of explicitly accounting for extrinsic and technical variability. To this end, we compared two differential-expression strategies: (i) a conventional analysis based on raw total RNA counts (*U* + *S*) using a two-fold change threshold, and (ii) a model-based approach in which mean expression was reconstructed from inferred kinetic parameters after correcting for the effective noise factor.

As shown in Fig. 6b, substantial discrepancies arise between the two strategies. Specifically, 53 genes identified as differentially expressed based on raw total-count means are reclassified as non-differential when reconstructed means are used, whereas 106 genes not detected by the raw-count approach are identified as differentially expressed under the model-based strategy. Representative examples of these two categories are shown in Figs. 6d and 6e (Supplementary Note 1).

In Fig. 6c, genes exhibiting significant fold changes in raw total-count means (|log_2_ FC| *>* 1) become non-significant when evaluated using reconstructed means, consistent with the absence of changes in inferred kinetic parameters such as burst size, burst frequency, or splicing time. Conversely, Fig. 6d shows genes with modest raw-count differences that become clearly separated based on reconstructed means, indicating that technical and extrinsic variability can obscure genuine biological differences at the count level. Kinetic analysis further provides mechanistic insight into these differences. For example, reduced reconstructed expression for genes such as *ITSN2* and *P2RY8* is primarily explained by decreases in burst size, whereas changes in the unspliced-to-spliced ratio reflect altered splicing time. In particular, the reduction of splicing time for *P2RY8* explains the observed decrease in the unspliced-to-spliced ratio. Together, these results demonstrate how PRIME disentangles mean-expression changes from their underlying kinetic regulation.

Finally, we examined known CTCL-associated markers, including *TRRAP*, which plays an important role in chromatin remodeling, and *RARA*, a key regulator of T-cell differentiation [72]. KDG analysis revealed a consistent kinetic signature characterized by increased burst frequency, prolonged splicing time, elevated total RNA abundance, and shifts in the unspliced-to-spliced ratio (Fig. 6e). Guided by this shared kinetic pattern, we identified three additional genes *PIK3R3, GATA3*, and *SLAMF1* (Fig. S6), all previously reported to be associated with CTCL [73–75]. Based on the same kinetic signature, we further identified *RSBN1* as a novel candidate marker gene displaying similar regulatory characteristics (Fig. 6e), suggesting a potential mechanistic link to CTCL progression.

## 3. DISCUSSION

We developed PRIME, a PGF-based framework that integrates mechanistic modeling with scalable inference to analyze single-cell RNA sequencing data using joint unspliced and spliced counts. By operating in PGF space, PRIME links stochastic gene-expression theory with practical single-cell analysis, enabling robust cell-state identification and mechanistically interpretable parameter inference within a unified framework.

A key insight of this work is that PGF-based representations provide a more stable and informative feature space than raw transcript counts. Because single-cell measurements are strongly affected by stochastic transcription, sampling variability, and technical noise, count-based representations can be unstable for downstream clustering. In contrast, PGFs encode distribution-level information and capture higher-order structure beyond mean expression. Across multiple datasets and clustering algorithms, we show that even a simple empirical PGF transformation substantially improves clustering performance, indicating that representation choice plays a central role in cell-state discovery.

PRIME further integrates a mechanistic transcription model that admits an exact steady-state PGF solution. This analytical tractability allows direct evaluation of model predictions without numerically solving the chemical master equation, enabling efficient genome-scale inference. By introducing a latent multiplicative factor to summarize cell-specific variability, PRIME accounts for confounded extrinsic and technical noise while preserving interpretable kinetic parameters. This provides a principled alternative to heuristic normalization procedures commonly used in single-cell analysis.

Compared with meK-means, a state-of-the-art biophysically guided clustering framework based on a negative-binomial-like description, PRIME is based on a more mechanistically explicit model in which the promoter switches between inactive and active states, transcription occurs from the active state, and the resulting RNA subsequently undergoes processing dynamics. This distinction is important because recent theoretical work [66] shows that negative-binomial-like behaviour can emerge across a broad region of parameter space and is therefore not uniquely indicative of transcriptional bursting. As a result, inference using a negative-binomial model may lead to strongly biased estimates of burst size and frequency. Such misspecification can in turn propagate to clustering errors, particularly when a subset of highly variable genes does not follow strongly bursty dynamics.

Algorithmically, PRIME combines a majorization-minimization clustering strategy with soft assignments and model-guided distances in PGF space. Kernel-density approximations over the latent variability factor replace explicit summation over cells, making computational cost largely independent of dataset size. Synthetic benchmarks and ablation analyses show that PGF representation, biophysical modeling, and soft assignment each contribute to performance gains, and together yield robust clustering and accurate parameter recovery.

Applications to experimental datasets demonstrate that PRIME achieves competitive or superior clustering accuracy compared with state-of-the-art methods while providing additional mechanistic insight. Importantly, PRIME enables identification of kinetically differentiated genes whose mean expression remains unchanged, highlighting regulatory differences that are invisible to conventional differential expression analysis. These results emphasize that transcriptional regulation is often encoded in kinetic parameters such as burst size, burst frequency, and processing time rather than in mean abundance alone.

Several limitations remain. PRIME currently assumes steady-state dynamics and a two-state promoter model, which does not capture transient dynamics nor does it explicitly model regulatory interactions such as the binding of transcription factors to promoters. In addition, PGF-based representations increase dimensionality, although this trade-off is offset by improved robustness and computational efficiency. Future work may extend the framework to time-resolved data, richer gene-state models, and additional multimodal measurements.

Overall, PRIME demonstrates that combining mechanistic modeling with PGF-based computation provides a powerful direction for single-cell analysis. By integrating probabilistic representation, scalable optimization, and Bayesian inference, the framework enables robust cell-state identification and genome-wide recovery of transcriptional kinetics, thereby advancing a more mechanistically grounded analysis of single-cell data.

## 5. ACKNOWLEDGMENTS

S. L., Y. W. and Q. J. acknowledge support from NSFC Grants (62573195 and 62322309), Shanghai Action Plan for Technological Innovation Grant (23S41900500). E. Z. C. acknowledges support from the Natural Science and Engineering Research Council of Canada’s (NSERC’s) Discovery Grant (RGPIN-2024-06015). R. G. acknowledges support from the Leverhulme Trust (RPG-2024-082).

## 6. METHODS

### Data sourcing and processing

We evaluated the performance of PRIME using five annotated scRNA-seq datasets spanning multiple tissues, species, and experimental designs. Two benchmark datasets were taken from the scRNA-seq mixology study [44], comprising mixtures of three human lung adenocarcinoma cell lines (HCC827, H1975 and H2228) and five human lung adenocarcinoma cell lines (HCC827, H1975, H2228, A549 and H838), denoted as `Cl3` and `Cl5`, respectively. A third annotated dataset, mouse primary motor cortex (denoted as `Mopsc`), was reported in Ref. [43]. For these three datasets, processed count matrices of spliced and unspliced RNAs together with curated cell-type annotations were directly sourced from the loom files released in [CaltechDATA repository](https://data.caltech.edu/records/2tsha-rcz19) [28].

In addition, we analyzed two annotated datasets processed from raw sequencing data. The `human skin` dataset comprises scRNA-seq profiles pooled from three healthy donors and sequenced using 10x Genomics v2 chemistry. The `mouse lung` dataset consists of lung mononuclear cells pooled from four mice under two conditions: *Streptococcus pneumoniae* infection and naïve controls. For both datasets, raw FASTQ files were obtained from the Gene Expression Omnibus (GEO; accession `GSE128066`) or alternatively downloaded from the BAMMSC GitHub repository [42]. FASTQ files were processed into BAM files using Cell Ranger v8.0.1 with the reference genomes GRCh38-2024-A and mm10-2020-A. Spliced and unspliced RNA count matrices were then generated using the `run10x` command in Velocyto v0.17.17 with the corresponding gene annotation files, and stored in loom format. Approximate cell-type annotations for the `human skin` and `mouse lung` datasets were obtained from [HumanSkin_ApproxTruth.RData](https://drive.google.com/drive/folders/1Dk4CLqSdkoR5E00zzV0wETlFZO1lCKLv) and [MouseLung_ApproxTruth.RData](https://drive.google.com/drive/folders/1ldz9MztRJgr2VvlKrzU7gOO4OaCBvgEH), respectively.

To assess PRIME’s capability for biological marker discovery in unlabeled settings, we further analyzed four unannotated scRNA-seq datasets. The `CTCL` dataset contains PBMCs collected from patients with cutaneous T-cell lymphoma and sequenced using ECCITE-seq [71]. As a control, the `ctrl` dataset consists of PBMCs collected from healthy donors. For both datasets, processed spliced and unspliced RNA count matrices were downloaded from Figshare as provided in Ref. [76]. The `Brca` dataset comprises scRNA-seq data from a mouse breast cancer model [70], with processed count matrices sourced from the [CaltechDATA repository](https://data.caltech.edu/records/n4yzp-qdf66) [28].

### Selection of highly variable genes

Highly variable genes were selected for each dataset following the protocol described in Ref. [28], which is based on the `highly_variable_genes` function implemented in Scanpy. The exact code used to identify HVGs for all datasets except `mouse lung` is publicly available in the GitHub repository [28]. For the `mouse lung` dataset, the precomputed list of HVGs was directly obtained from [MouseLung_ApproxTruth.RData](https://drive.google.com/drive/folders/1ldz9MztRJgr2VvlKrzU7gOO4OaCBvgEH), as provided in Ref. [42].

### Mechanistic model

In PRIME, we integrate spliced and unspliced RNA count data using a mechanistic transcription model with explicit gene-state switching and delayed splicing. The transcriptional dynamics of a single gene in a cell are described by

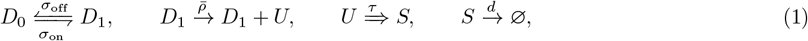

where the gene switches between an inactive state (*D*_0_) and an active state (*D*_1_) with rates *σ*_on_ and *σ*_off_. When the gene is active, unspliced transcripts *U* are produced at rate 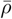; each unspliced transcript is converted into a spliced transcript *S* after a fixed processing delay *τ*; and spliced transcripts decay with rate *d*.

An exact steady-state solution for the joint distribution *P* (*U, S*) under Eq. (1) is available in PGF form [45]. Defining *u*_u_ = *z*_u_ − 1 and *u*_s_ = *z*_s_ − 1, the steady-state PGF 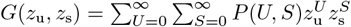 is

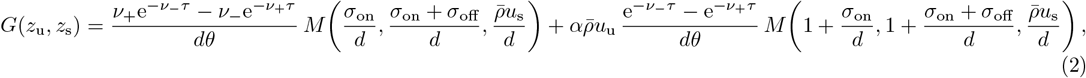

where *α* = *σ*_on_*/*(*σ*_on_ + *σ*_off_) and *M* (*a, b, x*) denotes the Kummer confluent hypergeometric function. The auxiliary variables are

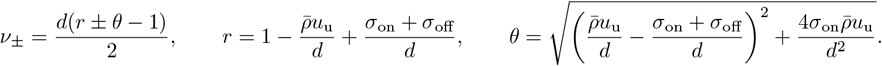

Following standard RNA velocity models [8, 77], we neglect degradation of unspliced RNA. In the regime *τd* ≪ 1, where the splicing delay is short relative to the lifetime of spliced RNA, Ref. [45] shows that the steady-state joint distribution generated by the delayed-splicing model in Eq. (1) is effectively indistinguishable from that obtained under a Markovian approximation, in which the original non-Markovian delay in splicing is replaced by a first-order reaction 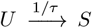 with an effective splicing rate 1*/τ*. We nevertheless retain the delayed formulation in Eq. (1) because it admits an exact closed-form PGF solution, whereas the corresponding first-order splicing model does not yield a closed-form expression for the steady-state joint distribution.

Equation (2) characterizes a homogeneous cell population in which all reaction-rate parameters in Eq. (1) are identical across cells, so that variability in (*U, S*) is dominated by intrinsic noise [78]. However, in reality, extrinsic noise due to a variation of parameters across the cell population, is also substantial. A major contributor to this type of noise is the coupling of transcriptional initiation to cell size (volume) [52, 79–81] which varies considerably from cell to cell [82]. To capture this effect, following recent extensions of the telegraph model [51, 83], we model the transcription (initiation) rate as

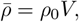

where *V* is the cell volume and *ρ*_0_ is a constant transcriptional activity per unit volume. Note that transcription maybe dependent on the nuclear volume rather than the cellular volume but since these are typically proportional to each other [84] this distinction is not relevant to our model.

We additionally account for technical noise arising from incomplete transcript capture (dropout) in scRNA-seq. We use a binomial capture model [54], in which each transcript (unspliced or spliced) is independently captured and sequenced with probability *p*_cap_, which may vary across cells. As shown in Supplementary Note 2, this capture process induces a simple rescaling of the PGF variables and is mathematically equivalent to the effective transformation

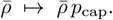

Consequently, in the absence of additional orthogonal measurements, volume-driven extrinsic variability (via *V*) and capture-related technical variability (via *p*_cap_) are statistically confounded, because both enter the model only through the same multiplicative scaling of the effective transcription rate 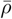 [66]. We therefore introduce a single latent effective factor *β* to summarize these cell-to-cell effects and write

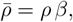

where *ρ* is an effective rate coefficient and *β* captures the combined contribution of volume and capture efficiency. In practice, the distribution of *β* can be estimated from the normalized per-cell total transcript abundance (i.e., the library-size-adjusted total counts), which provides a direct proxy for this multiplicative variability across cells (Eq. (S6) in Supplementary Note 2).

Since Eq. (2) describes the steady-state solution, the five kinetic parameters in the model Eq. (1) are not independently identifiable. Instead, only their ratios relative to the degradation rate can be uniquely determined, namely (*ρ/d, σ*_on_*/d, σ*_off_*/d, τd*). Therefore, without loss of generality, we set *d* = 1 throughout this paper, which renders the remaining four kinetic parameters dimensionless.

### PGF-based majorization-minimization clustering method

Let 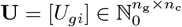 and 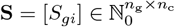 denote the count matrices of unspliced and spliced RNAs, where *n*_g_ and *n*_c_ are the numbers of genes and cells, respectively. We denote the full gene set by 𝔾_ent_ and the set of HVGs by 𝔾_hvg_, with *n*_h_ = |𝔾_hvg_|.

To account for cell-specific technical and extrinsic variability, we use the effective noise factor (*β*_*i*_) for cell *i* that rescales the transcription rate in the mechanistic model, which can be estimated by

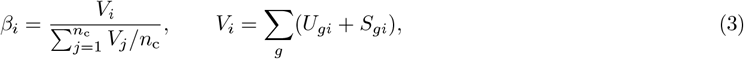

For some HVG *g*, the empirical PGF of cell *i* is

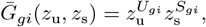

while the model PGF under cluster *k* is given analytically as *G*(*z*_u_, *z*_s_ | *φ*_*gk*_, *β*_*i*_), where *φ*_*gk*_ denotes the kinetic parameters {*ρ, σ*_on_, *σ*_off_, *τ*}.

We measure the discrepancy between a cell and a cluster in PGF space by

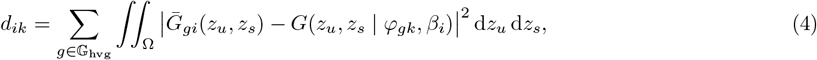

where Ω = [*z*_min_, *z*_max_]^2^ with *z*_min_ = 0.95 and *z*_max_ = 1. The integrals are evaluated using Gauss-Legendre quadrature.

To obtain robust assignments under single-cell sampling noise, we aggregate distances using the power (Kolmogorov) mean with exponent *s <* 0,

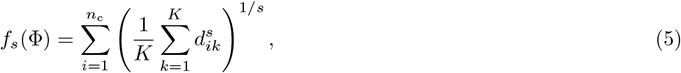

where Φ denotes the collection of kinetic parameters *φ*_*gk*_ for *g* ∈ 𝔾_hvg_ and *k* = 1, · · ·, *K*, and *K* is the total number of clusters. As *s* → − ∞, this recovers the hard minimum underlying *K*-means; finite negative *s* performs soft pooling across clusters and mitigates sensitivity to noise.

The clustering problem can thus be formulated as an optimization problem in which we seek Φ that minimizes *f*_*s*_(Φ). To solve this problem, we adopt a majorization-minimization (MM) strategy. Because the power-mean objective *f*_*s*_(Φ) is concave in *d*_*ik*_, we construct a first-order upper bound at the current parameter estimate Φ^(*m*)^ at iteration *m*. This yields the surrogate objective

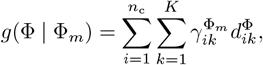

where the responsibilities

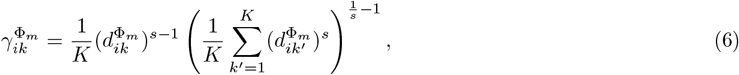

act as soft cluster assignments. This majorization step effectively computes the responsibilities, analogous to the expectation step in the expectation-maximization (EM) algorithm. The subsequent minimization step solves

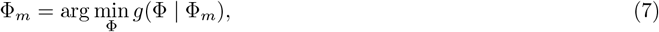

which corresponds to the maximization step in EM. Direct evaluation of the surrogate objective *g*(Φ | Φ_*m*_) requires summation over all values of the effective noise factor *β*, causing the computational cost to scale with the number of cells. To mitigate this, we approximate these sums using kernel density estimation (KDE) over *β* and evaluate the resulting integrals via Gaussian-Legendre quadrature. This reduces the complexity of each update to depend only on the quadrature resolution rather than *n*_c_. Moreover, the surrogate objective *g*(Φ | Φ_*m*_) is separable across gene-cluster pairs (*g, k*), enabling straightforward parallelization. Each resulting subproblem is solved using the derivative-free Nelder-Mead method implemented in `Optim.jl`. Once a majorization-minimization cycle is completed, the power parameter *s* is scaled by a factor *η* = 1.5, which progressively sharpens the soft assignments and ensures that the final solution approaches a hard clustering configuration.

A full derivation is provided in Supplementary Note 3, where the complete procedure is summarized in Algorithm 1.

### PGF-based Bayesian parameter inference

After convergence of the majorization-minimization procedure, cells are assigned to clusters {𝒞_*k*_} according to their responsibilities. Specifically, cell *i* is assigned to the cluster

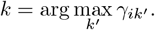

For each cluster *k*, we define 𝒞_*k*_ ⊆ {1, …, *n*_c_} as the set of cells whose maximum responsibility occurs at *k*, and denote its size by |𝒞_*k*_|.

For any given gene *g* ∈ 𝔾_ent_ and cell cluster 𝒞_*k*_, we define the empirical joint PGF of unspliced and spliced counts as

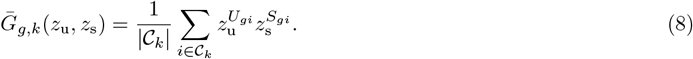

To account for multiplicative cell-to-cell variability, we associate each cell *i* with an effective noise factor *β*_*i*_ *>* 0. For cluster *k*, we estimate the distribution of *β* using KDE,

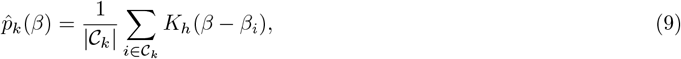

where *K*_*h*_ is a kernel with bandwidth *h*. Let *φ* = {*ρ, σ*_on_, *σ*_off_, *τ*} denote the kinetic parameters to be inferred. The model-implied PGF for gene *g* in cluster *k* is then given by the *β*-mixture

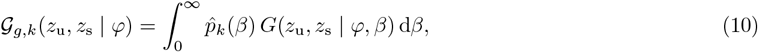

where *G*(*z*_u_, *z*_s_ | *φ, β*) is obtained from Eq. (2) by substituting 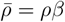.

We evaluate the integral in Eq. (10) using Gauss-Legendre quadrature over the interval [*β*_min_, *β*_max_], where 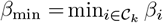 and 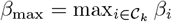. Specifically,

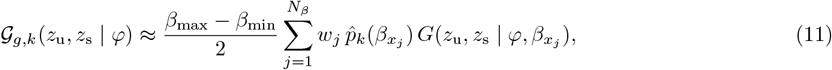

with

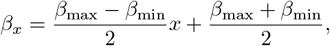

where 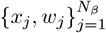 are the quadrature nodes and weights computed using the `gausslegendre` function in Julia.

We sample the PGF on a fixed grid by selecting *n*_*z*_ = 7 evenly spaced values for each of *z*_u_ and *z*_s_ within the interval [0.95, 0.99]. This choice matches exactly the *z*_u_ and *z*_s_ range and resolution used during clustering. The resulting Cartesian product forms 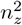 evaluation points,

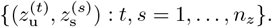

Evaluating both the empirical and model PGFs on this grid yields the vectors

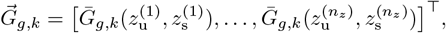

and

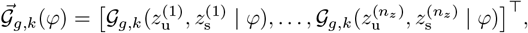

with identical ordering in both vectors.

As shown in Supplementary Note 4, if the empirical PGF is constructed from samples drawn from the distribution encoded by 𝒢_*g,k*_(·, · | *φ*^∗^), where *φ*^∗^ denotes the ground-truth parameters, then

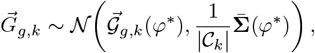

with covariance entries

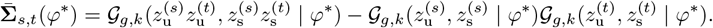

A direct Bayesian likelihood would therefore be

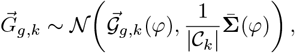

but the dependence of 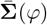 on the parameters *φ* makes inference numerically unstable. To ensure computational tractability, we replace 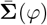 with an empirical plug-in covariance computed directly from 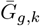,

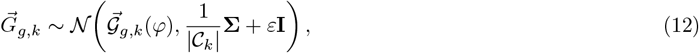

where

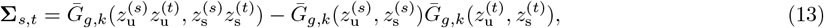

and *ε* is a very small number to prevent numerical singularity. Posterior inference for *φ* = {*ρ, σ*_on_, *σ*_off_, *τ*} was performed in Julia using `Turing.jl` [85]. We adopted weakly informative priors,

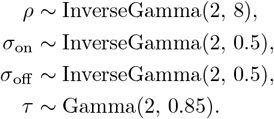

These priors provide broad support for the transcription rate *ρ*, allowing orders-of-magnitude variation in expression levels, while constraining the switching rates *σ*_on_ and *σ*_off_ to biologically plausible values on the order of tens and *τ* to moderate values. These ranges are consistent with empirical measurements reported in Ref. [83]. The full PGF-based Bayesian parameter inference procedure is summarized in Algorithm 2 in Supplementary Note 3.

### Identification of kinetically differentiated genes (KDGs)

The identification of kinetically differentiated genes proceeds through four stages: (i) data-level quality control prior to parameter inference; (ii) PGF-based Bayesian inference of kinetic parameters; (iii) post-inference quality control of parameter estimates; and (iv) statistical testing for kinetic differentiation between clusters.

#### (i) Data-level quality control

After clustering with PRIME, we first remove extreme outliers to ensure robust parameter estimation. For each gene *g* ∈ 𝔾_ent_ and cluster *k*, only cells *i* ∈ 𝒞_*k*_ satisfying

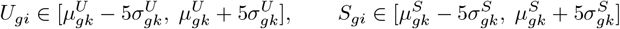

are retained for inference. Here,

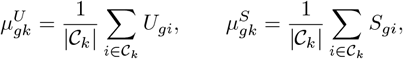

and

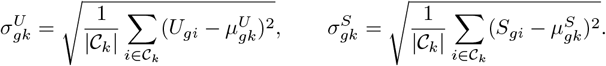

Cells outside these ranges are excluded to prevent rare extreme observations from disproportionately influencing posterior inference.

#### (ii) PGF-based Bayesian parameter inference

For each gene *g* in each cluster *k*, we perform Bayesian inference under the steady-state assumption of the mechanistic model (Eq. (1)). Without loss of generality, we fix *d* = 1, so that inferred parameters are normalized relative to the degradation rate. Posterior samples are obtained for the four kinetic parameters

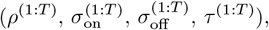

where *T* denotes the total number of MCMC samples. From these traces, we compute posterior samples of three dimensionless quantities:

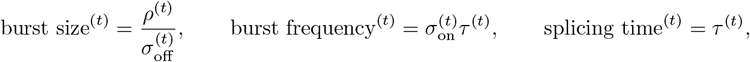

for *t* = 1, …, *T*. Point estimates are defined as posterior medians, denoted med(*x*).

#### (iii) Quality control of inferred parameters

To assess estimation reliability, we compute the highest density interval (HDI) with 80% posterior probability. Given ordered samples *x*_1_ ≤ · · · ≤ *x*_*T*_ and Δ = ⌊0.8*T* ⌋, we identify

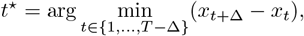

and define the interval width

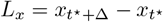

computed by the built-in function `hdp` in Julia. We quantify relative precision via

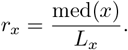

Additionally, MCMC sampling quality is evaluated using the effective sample size (ESS) computed by the built-in function `ess` in Julia. A parameter estimate passes quality control if

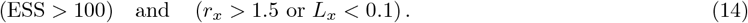

The auxiliary criterion *L*_*x*_ *<* 0.1 prevents rejection of small but precisely estimated parameters for which *r*_*x*_ may be artificially low. Only parameters satisfying Eq. (14) are retained for downstream KDG analysis.

#### (iv) Statistical identification of KDGs

For a gene *g* and two clusters *a* and *b*, the fold change (FC) of a kinetic parameter *x* is defined as

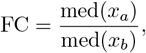

and log_2_ FC serves as the horizontal axis in volcano plots.

Statistical significance is assessed using the posterior superiority probability (PS),

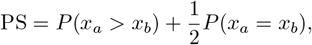

with two-sided *p*-value

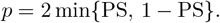

The value log_10_ *p* serves as the vertical axis in volcano plots. In practice, PS is approximated using posterior samples:

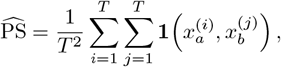

where **1** equals 1, 0.5, or 0 depending on whether 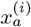 is greater than, equal to, or less than 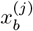.

A gene is declared kinetically differentiated for parameter *x* if

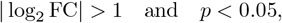

together with the mean-expression constraint

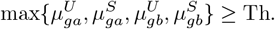

This additional threshold ensures that differentiation is not driven by extremely low-expression genes dominated by technical noise. The value of Th is dataset-specific: for the `Brca` dataset, Th = 1, whereas for the `CTCL` and `ctrl` datasets, Th = 0.1.

### Identification of differentially expressed genes (DEGs)

For a gene *g* ∈ 𝔾_ent_ and two clusters *a* and *b*, differential expression is assessed based on the total transcript abundance, defined as the sum of unspliced and spliced counts. The mean expression fold change is computed as

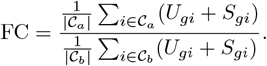

Statistical significance *p*-value is evaluated using a two-sided Mann-Whitney U test applied to the per-cell total counts

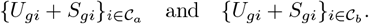

A gene is classified as differentially expressed if it satisfies both

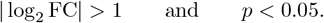

## Supplementary Materials

### Supplementary Note 1

#### Fig. 1

- K-means: Clustering was performed using the Python packages `Scanpy` and `scikit-learn`. We applied `KMeans` to the feature matrix `adata.X`, with the number of clusters fixed to the number of ground-truth labels.
- Leiden: Leiden clustering was implemented in `Scanpy`. A neighborhood graph was first constructed using `sc.pp.neighbors(adata, use rep=“X”)`, followed by community detection via `sc.tl.leiden` with resolution set to 1 and otherwise default parameters.
- Fuzzy C-means: Fuzzy clustering was carried out using the Julia package `Clustering.jl`. We used `fuzzy_cmeans(X, k, 1.2, maxiter=10000)`, where *k* denotes the target number of clusters and the fuzziness coefficient was set to 1.2.
- t-SNE: Two-dimensional embeddings were generated in R using `Rtsne`, with PCA initialization enabled, perplexity set to 20, and the maximum number of iterations set to 1000.

#### Fig. 3

##### Synthetic data generation

For each cell *i*, we first sampled an effective noise factor

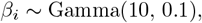

which rescales the transcription rate. We considered two experimental settings. *Case 1: K* = 5 clusters with *n*_g_ = 600 genes. The numbers of cells per cluster were 200, 400, 800, 1600, and 2000. *Case 2: K* = 10 clusters with *n*_g_ = 1100 genes. The numbers of cells per cluster were 100, 100, 300, 300, 500, 500, 800, 1000, 1000, and 1400. In both cases, the number of HVGs was fixed at *n*_h_ = 100. For genes *g* ∈ 𝔾_ent_ \ 𝔾_hvg_, parameters in Eq. (1) were sampled independently from lognormal distributions [29, 86] as

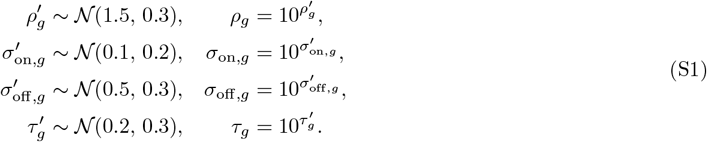

These parameters were shared across clusters. For *g* ∈ 𝔾_hvg_, we introduced differential expression by sampling, for each cluster *k*,

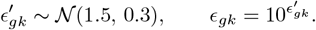

The parameters *σ*_on,*g*_, *σ*_off,*g*_, and *τ*_*g*_ were drawn from Eq. (S1) as above, while the effective transcription rate in cluster *k* was defined as

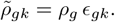

For each cell *i* and gene *g*, the count pair (*U*_*gi*_, *S*_*gi*_) in steady-state conditions was generated by simulating the delayed transcription model in Eq. (1) using `DelaySSAToolkit.jl` [57] with degradation rate *d* = 1. For non-HVGs, simulations used parameters {*ρ*_*g*_*β*_*i*_, *σ*_on,*g*_, *σ*_off,*g*_, *τ*_*g*_}. For HVGs in a cell belonging to cluster *k*, we replaced *ρ*_*g*_ by 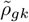, resulting in 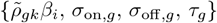.

##### Ablation study (Fig. 3a)

To dissect the contribution of each component of PRIME, we constructed several reduced variants by selectively removing key ingredients of the full framework. The variants are summarized below.

- **Reduce 1** (PGF representation + noise correction + biophysical model): After each parameter update, hard cluster assignments are enforced instead of soft responsibilities. Given the inferred kinetic parameters at iteration *m*, the cell-cluster distance 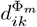 is computed using Eq. (S13), and cell *i* is assigned to the nearest cluster,

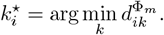

Responsibilities are then replaced by hard indicators,

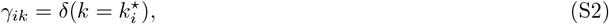

where *δ*(·) denotes the Kronecker delta. The minimization step is subsequently performed using Eq. (S18) together with Eq. (S2). This variant isolates the effect of soft assignment together with power K-means in PRIME.
- **Reduce 2** (PGF representation + biophysical model): This variant follows Reduce 1 but removes correction for extrinsic and technical variability by fixing *β*_*i*_ = 1 for all cells. It therefore evaluates the contribution of modeling cell-specific multiplicative noise.
- **Reduce 3** (PGF representation + soft cluster assignment + power K-means backbone): This method retains the PGF representation and the power-mean objective but removes the biophysical model. In Eq. (S11), the model PGF is replaced by a cluster centroid in PGF space,

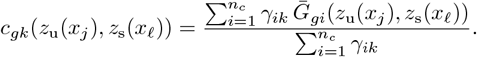

Responsibilities are updated according to Eq. (S17). Since no kinetic model is used, *γ*_*ik*_ is independent of model parameters.
- **Reduce 4** (PGF representation): Cells are represented using PGF features as in Fig. 1, followed by standard K-means clustering. This isolates the contribution of PGF representation alone without model-based inference or power K-means backbone.
- **Reduce 5** (Count representation + soft cluster assignment + power K-means backbone): For each cell, unspliced and spliced counts of HVGs are concatenated into a feature vector

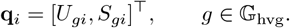

Cluster distances are computed using Euclidean distance,

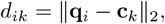

where the centroid is defined as

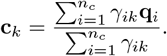

Responsibilities are updated using Eq. (S17). This variant evaluates the effect of replacing PGF features with raw counts while keeping soft cluster assignment and power K-means backbone.
- **Reduce 6** (Count representation + noise correction + soft cluster assignment + power K-means backbone): This variant first normalizes counts by the effective noise factor estimated from Eqs. (S7) and (S8),

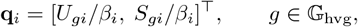

and then follows the same procedure as Reduce 5. It also evaluates whether simple normalization can substitute explicit modeling of *β*.
- **Reduce 7** (Count representation): Unspliced and spliced counts are concatenated into **q**_*i*_ = [*U*_*gi*_, *S*_*gi*_]^⊤^, and standard K-means clustering is directly applied in count space.
- **Reduce 8** (Count representation + noise correction): Counts are first normalized by *β*_*i*_,

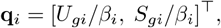

and standard K-means clustering is then performed. This variant evaluates the benefit of simple normalization without PGF transformation or soft assignment.

##### Comparative study (Fig. 3b)

To ensure a fair comparison, all methods were applied to the same synthetic datasets using identical input matrices of unspliced and spliced RNA counts. Unless otherwise specified, the number of clusters *K* was set to the ground-truth value used in data generation. Below we summarize the implementation details of each competing method.

- **meK-means**: We followed the official implementation and protocol provided by the authors of meK-means (https://github.com/pachterlab/CGP_2023/blob/main/example_meK-means_notebook.ipynb). In addition to unspliced and spliced count matrices, meK-means requires gene-length information. Because gene length is not explicitly modeled in our synthetic data generation, all genes were assigned a constant length of 10^5^ for consistency. The number of clusters was set to the ground-truth value, and all other parameters were kept at their default settings.
- **log+K-means**: Normalization and logarithmic transformation were performed using Scanpy commands `sc.pp. normalize total` followed by `sc.pp.log1p`. K-means clustering was then applied to the transformed feature matrix using the same protocol described for Fig. 1.
- **log+Leiden**: The same normalization and log-transformation procedure as in log+K-means was applied. A nearest-neighbor graph was constructed using `sc.pp.neighbors`, followed by Leiden clustering via `sc.tl.leiden`, consistent with the setup used in Fig. 1.
- **Scranpooling+K-means**: Normalization was performed in R using the `scran` package following the official vignette (https://bioconductor.org/packages/release/bioc/vignettes/scran/inst/doc/scran.html). Specifically, `quickCluster`, `computeSumFactors`, and `logNormCounts` were applied sequentially. The normalized data were then exported to Python, and K-means clustering was performed following the protocol used in Fig. 1.
- **Scranpooling+Leiden**: The same Scran normalization pipeline as above was applied. After importing the normalized data into Python, Leiden clustering was performed using the standard Scanpy workflow described for Fig. 1.
- **scVI+K-means**: We used the Python package `scvi-tools` following the official documentation (https://scvi-tools.org/). Unspliced and spliced counts were treated as two modalities within the model. After training (`model.train`), the latent embedding was extracted using `model.get_latent_representation`, and K-means clustering was applied to this latent space.
- **scVI+Leiden**: The same scVI pipeline as above was used to obtain the latent representation. Leiden clustering was then performed on the latent embedding following the standard Scanpy workflow.
- **WNN+Leiden**: Weighted Nearest Neighbor (WNN) integration was implemented using `pyWNN` (https://github.com/dylkot/pyWNN). Spliced and unspliced matrices were normalized using `normalize_total` and `log1p`, followed by PCA independently for each modality (20 principal components retained). A WNN graph was constructed with `n_neighbors=20`, and Leiden clustering was performed on this graph using `sc.tl.leiden` with `neighbors_key=“WNN”` and `resolution=1`.
- **MOFA2+K-means**: We used the Python package `mofapy2` following the official tutorial (https://github.com/bioFAM/mofapy2/blob/master/mofapy2/notebooks/getting_started_python.ipynb). Unspliced and spliced counts were treated as two separate views to construct a `MuData` object. Latent factors learned by MOFA2 were extracted and used as input features for K-means clustering.
- **MOFA2+Leiden**: The same MOFA2 workflow was used to infer latent factors, followed by Leiden clustering applied to the learned factor representation.
- **MAGIC+K-means**: We used the Python package `magic-impute` (`magic.py`) following the official tutorial (https://nbviewer.org/github/KrishnaswamyLab/MAGIC/blob/master/python/tutorial_notebooks/emt_tutorial.ipynb). Unspliced and spliced count matrices were concatenated, and data smoothing was performed using `MAGIC().fit_transform`. The resulting imputed features were then clustered using K-means.
- **MAGIC+Leiden**: The same MAGIC preprocessing was applied, and Leiden clustering was performed on the smoothed representation using the Scanpy workflow.
- **DIMM-SC**: DIMM-SC employs a hierarchical Dirichlet mixture model. We followed the official implementation provided in https://github.com/CHPGenetics/BAMMSC. Clustering was performed using the command `BAMMSC` with the configuration `option=“DIMMSC”`.

##### Parameter inference & computational efficiency (Fig. 3c–3e)

We compare PRIME with meK-means under matched model assumptions, where the meK-means model is aligned with the model used for data generation. To ensure a fair comparison, particularly in terms of computational efficiency, we implemented meK-means in Julia. In this modified meK-means, the joint probability distribution of model Eq. (1) is computed using the finite state projection (FSP) method [63], with the resulting system of ordinary differential equations solved using `DifferentialEquations.jl`. In practice, this corresponds to solving the finite system defined by Eqs. (S46), (S49), and (S51) in the Supplementary of Ref. [45]. The original meK-means implementation converts PGF solutions into joint probabilities using an inverse fast Fourier transform (IFFT). However, because the analytical PGF solution (Eq. (2)) of Eq. (1) involves hypergeometric functions, our tests show that computing probabilities via IFFT is substantially slower than using the FSP approach. In the M-step of meK-means (corresponding to the minimization step in PRIME), we retain the original Kullback–Leibler divergence objective and solve the resulting optimization problem using the same optimizer as PRIME, namely the Nelder–Mead method. For both methods, the convergence tolerance is set to `g_tol=`10^−9^. The maximum number of iterations is set to `maxiter=200` for meK-means and `maxiter=900` for PRIME, placing PRIME at a disadvantage in the computational efficiency comparison. PRIME performs 10 majorization–minimization iterations, whereas meK-means performs 10 EM iterations.

##### Computational efficiency of parameter inference (Fig. 3f)

We compare PRIME’s Bayesian parameter inference method with a count-based Bayesian estimator whose likelihood is computed using the FSP approach. In the latter, the joint probability distribution is obtained by solving the system of ODEs defined by Eqs. (S46), (S49), and (S51) in the Supplementary of Ref. [45]. Both methods are applied to the first 10 genes in a cluster of 1000 cells from the *K* = 5 synthetic dataset. The ground-truth parameter values for the 10 genes are listed in Table S1. Posterior sampling is performed using the `NUTS` sampler with 1000 iterations for both methods.

#### Fig. 4

*Fig. 4a*. Because the `human skin` and `mouse lung` datasets contain explicit batch structure (multiple donors or experimental conditions), we additionally compared PRIME with clustering pipelines that incorporate dedicated batch-effect correction methods. Specifically, we evaluated ComBat- and MNN-based corrections, both of which require predefined batch labels.

- **ComBat+K-means**. Batch-effect correction was performed using the Scanpy implementation of ComBat via `scanpy.pp.combat`, with batch labels provided according to the dataset metadata. The corrected expression matrix was then used as input for K-means clustering following the same protocol described for Fig. 1.
- **ComBat+Leiden**. ComBat correction was applied using `scanpy.pp.combat` as above. A nearest-neighbor graph was subsequently constructed from the corrected representation, and Leiden clustering was performed using the standard Scanpy workflow.
- **MNN+K-means**. Batch correction was carried out using the Python package `mnnpy` via the function `mnn correct`, with cells grouped by batch labels. The corrected feature matrix was then clustered using K-means under the same settings as in Fig. 1.
- **MNN+Leiden**. After applying MNN correction using `mnnpy::mnn_correct`, Leiden clustering was performed on the corrected representation following the standard Scanpy pipeline.
- **MeK-means**. For the experimental datasets `cl3`, `cl5`, and `Mopsc`, we used the published gene-length reference files released alongside Ref. [28] and available at https://github.com/pachterlab/CGP_2023/tree/main/references. For `human skin` and `mouse lung`, gene-length reference files were generated following the reference-construction procedure described at https://github.com/pachterlab/GP_2021_3/tree/main/processing_scripts/make_references.

*Fig. 4b*. To visualize the clustering results, we performed dimensionality reduction using t-distributed stochastic neighbor embedding (t-SNE). Specifically, t-SNE was implemented in R using the `Rtsne` package. The input features were first projected using principal component analysis (`pca=TRUE`), and t-SNE was then run with `perplexity=20` and `max_iter=1000`. The resulting two-dimensional embedding was used solely for visualization.

*Fig. 4c*. To align inferred clusters with known cell-type annotations, we evaluated all possible permutations between predicted cluster labels and reference labels, and selected the mapping that maximized the ARI. The resulting optimal permutation was used to define the aligned correspondence between inferred clusters and annotated cell types.

#### Fig. 6

*Fig. 6a*. Cell types were annotated using cell-surface protein markers reported in Table S2 of the Supplementary Materials of Ref. [76]. Specifically, CD3 together with CD4 was used to identify CD4^+^ T cells, CD3 together with CD8 was used to identify CD8^+^ T cells, and CD16 was used to annotate monocytes. After clustering, the average abundance of each marker protein was computed within each cluster and standardized using *z*-score normalization across clusters. Clusters were then aligned to cell types based on their normalized marker-enrichment profiles.

*Fig. 6d,e*. Fold-change distributions of kinetic parameters (burst size, burst frequency, splicing time), normalized total counts, and count ratios were estimated using a bootstrap-like cross-validation procedure. For each sample, cells were randomly partitioned into 10 folds. In each run, one fold was held out and PRIME’s PGF-based parameter inference was performed using the remaining nine folds to obtain point estimates. Repeating this procedure generated 10 estimates for each parameter in each cluster. Fold-change distributions were then constructed by computing all pairwise combinations (10 × 10) between the two samples.

### Supplementary Note 2

Consider a cell of volume *V* and a particular gene of interest. The joint distribution of unspliced and spliced RNA counts is encoded by the probability generating function (PGF), whose closed-form expression is given in Eq. (2). Inspecting Eq. (2) shows that the transcription rate 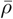 and the shifted PGF variables (*u*_u_ = *z*_u_ − 1, *u*_s_ = *z*_s_ − 1) always enter as paired combinations.

We now incorporate the capture (dropout) process. Let random variables *U* and *S* denote the ground-truth (pre-capture) counts of unspliced and spliced transcripts in a cell. Conditional on *V*, the PGF of their joint distribution is

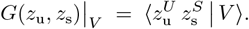

We assume that each unspliced transcript is captured and sequenced independently with probability *p*_cap_, and likewise for each spliced transcript (with the same capture probability). Introduce Bernoulli random variables

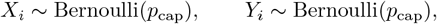

so that *P* (*X*_*i*_ = 1) = *P* (*Y*_*i*_ = 1) = *p*_cap_ and *P* (*X*_*i*_ = 0) = *P* (*Y*_*i*_ = 0) = 1 − *p*_cap_. The observed counts are then

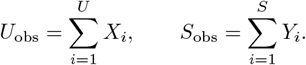

Given (*V, p*_cap_), the observed PGF is

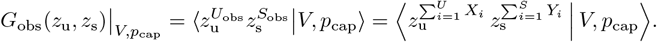

By the law of total expectation, conditioning on (*U, S*) yields

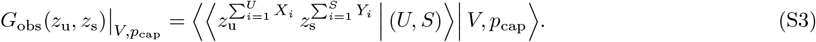

Fix (*U, S*) = (*m, n*). Then 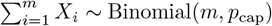 and 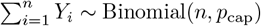. Since

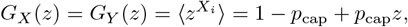

and 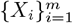 are i.i.d., 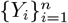 are i.i.d., and the two families are mutually independent, we obtain

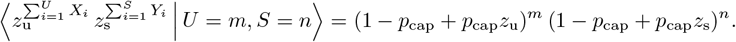

Substituting this expression into Eq. (S3) and reintroducing (*U, S*) as random variables gives

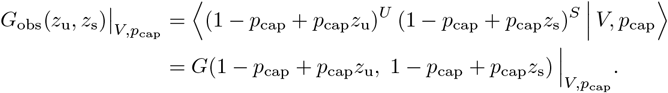

Therefore, the binomial capture model induces a simple affine mapping of the PGF arguments [56]

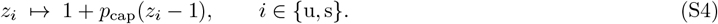

Equivalently, in shifted variables *u*_u_ = *z*_u_ − 1 and *u*_s_ = *z*_s_ − 1, Eq. (S4) becomes a multiplicative scaling,

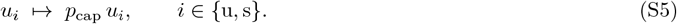

Finally, because 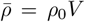 and Eq. (2) depends on the shifted PGF variables only through the paired products 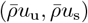, the binomial capture model in Eq. (S5) implies that capture enters the observed PGF through the same multiplicative structure. In particular, for fixed (*V, p*_cap_) we have the mapping

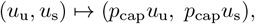

so that 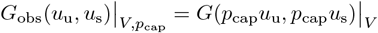. For a heterogeneous population with variability in both cell volume *V* and capture probability *p*_cap_, the marginal observed PGF is therefore obtained by averaging over their distributions *p*(*V*) and *p*(*p*_cap_):

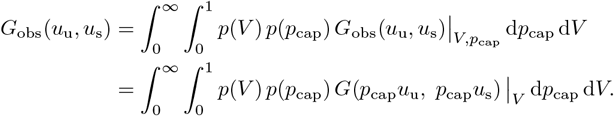

This structure shows that volume-driven extrinsic variability and capture-related technical variability enter only through the product *V p*_cap_. We therefore introduce an effective latent variable

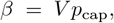

and rewrite the effective transcription rate as 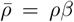. With this reparameterization, the observed PGF can be expressed as a one-dimensional mixture,

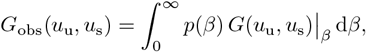

where *G*(*u*_u_, *u*_s_) | _*β*_ denotes Eq. (2) evaluated at 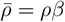.

Differentiating *G*(*u*_u_, *u*_s_)|_*β*_ with respect to *u*_u_ and *u*_s_ and evaluating at (*u*_u_, *u*_s_) = (0, 0) yields the conditional means

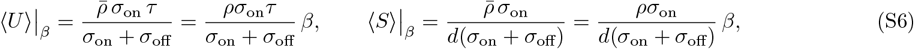

and hence ⟨*U* + *S*⟩ |_*β*_ ∝ *β*. This motivates estimating *p*(*β*) directly from cell-wise total transcript abundance (after normalization), thereby jointly accounting for extrinsic (volume) and technical (capture) variability through a single effective factor.

### Supplementary Note 3

Let 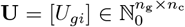 and 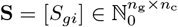 denote the count matrices of unspliced and spliced RNAs, where *n*_g_ and *n*_c_ are the number of genes and cells, respectively. We denote the full gene set by 𝔾_ent_ and the set of highly variable genes (HVGs) by 𝔾_hvg_, with *n*_h_ = |𝔾_hvg_|. Let *K* denote the number of cell clusters.

For each cell *i*, we define an effective noise factor *β*_*i*_ to reflect both extrinsic and technical variability:

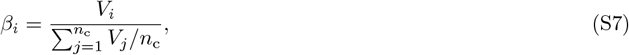

where

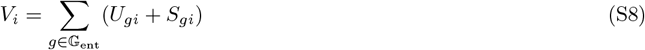

is the total number of transcripts observed in cell *i*.

For each gene *g* ∈ 𝔾_hvg_ and cell *i*, the empirical PGF is given by

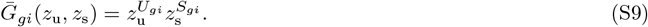

We model the mechanistic transcription dynamics using Eq. (1), where the effective transcription rate in cell *i* is 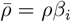. The parameter set for gene *g* in cluster *k* is denoted by *φ*_*gk*_ = {*ρ, σ*_on_, *σ*_off_, *τ*}. The model PGF for gene *g* in cell *i* under cluster *k* is then *G*(*z*_u_, *z*_s_ | *φ*_*gk*_, *β*_*i*_) computed via the analytical form in Eq. (2).

We define the PGF distance between cell *i* and cluster *k* for gene *g* as

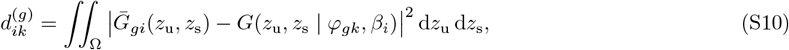

where the integration domain is Ω = [*z*_min_, *z*_max_]^2^, with *z*_min_ = 0.95 and *z*_max_ = 1. These limits are chosen to improve discriminability between cell populations in the PGF space, as shown in [87].

To accelerate computation of Eq. (S10), we apply 2D Gauss–Legendre quadrature:

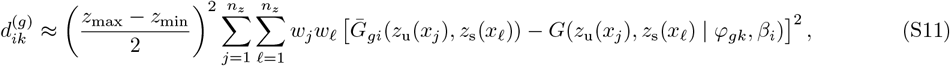

where

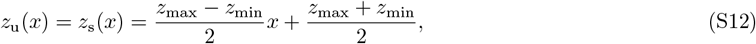

and 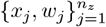 are the nodes and weights of the quadrature, computed using the `gausslegendre` function in Julia. The overall PGF-based distance from cell *i* to cluster *k* is aggregated over HVGs:

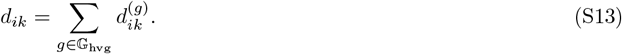

To define the clustering loss, we introduce the *Kolmogorov mean*. For a vector **x** = [*x*_1_, …, *x*_*n*_]^⊤^ with *x*_*i*_ *>* 0 and a strictly monotonic function *g*: ℝ_*>*0_ → ℝ, the Kolmogorov mean is defined as

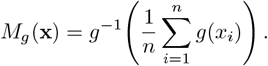

In this work, we focus on the family of power means with *g*(*x*) = *x*^*s*^ for *s <* 0, and denote the resulting mean by *M*_*s*_(**x**).

Let **d**_*i*_ = [*d*_*i*1_, …, *d*_*iK*_]^⊤^ denote the PGF-based distances from cell *i* to the *K* clusters. We define the clustering loss as

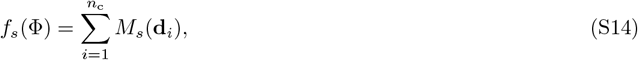

where Φ = {*φ*_*gk*_} for all *g* ∈ 𝔾_hvg_ and *k* = 1, …, *K*.

As *s* → −∞, the power mean satisfies *M*_*s*_(**d**_*i*_) → min_*k*_ *d*_*ik*_, and the loss in Eq. (S14) reduces to the classical *K*-means objective [41]. However, in our setting, each empirical PGF (Eq. (S9)) is constructed from a single observation. Consequently, the resulting empirical distribution can be highly variable and may deviate substantially from the underlying cluster-level PGF, particularly under limited sample size. As demonstrated in Fig. 3a (Reduce 1 and Reduce 2), such variability can cause the minimum-distance assignment to be unstable, leading to misclassification when sampling noise dominates true cluster structure. To mitigate this effect, we adopt power means with moderate negative *s*, which softly aggregate distances across clusters and effectively perform partial pooling. This weighting scheme reduces sensitivity to spurious minima and improves clustering robustness in low-sample regimes.

The optimization problem in Eq. (S14) is nontrivial. We therefore adopt a majorization-minimization (MM) strategy, which generalizes the expectation-maximization (EM) framework [88]. An MM algorithm iteratively minimizes a sequence of surrogate functions *g*(Φ | Φ_*m*_) that majorize the objective *f*_*s*_(Φ) at the current iterate Φ_*m*_. Specifically, the surrogate satisfies the tangency condition *g*(Φ_*m*_ | Φ_*m*_) = *f*_*s*_(Φ_*m*_) and the domination condition *g*(Φ | Φ_*m*_) ≥ *f*_*s*_(Φ) for all Φ. These properties ensure the descent relation

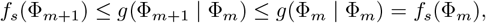

where the second inequality follows from minimizing *g*(Φ | Φ_*m*_) with respect to Φ. Repeated application of this inequality guarantees monotonic convergence of the MM algorithm.

To construct the surrogate, we first compute the derivative of *M*_*s*_(**d**_*i*_) with respect to *d*_*ik*_,

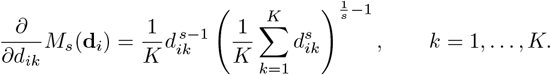

To make the dependence on model parameters explicit, we write 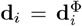. For *s <* 1, the function 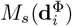 is concave in 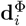. Therefore, for a given iterate Φ_*m*_, it admits the first-order upper bound

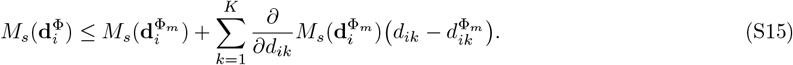

Summing Eq. (S15) over all cells yields the majorization of the loss function,

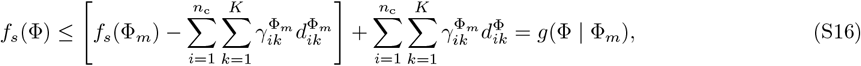

where

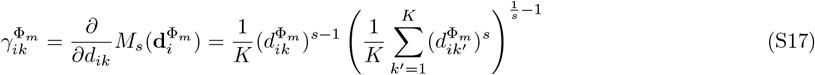

are the responsibility weights, which quantify the soft assignment probability of cell *i* to cluster *k*. This step is analogous to the expectation step of the classical EM algorithm.

Here, the first term on the right-hand side of Eq. (S16) is constant because Φ_*m*_ is fixed at iteration *m*. Consequently, minimizing *g*(Φ | Φ_*m*_) is equivalent to solving

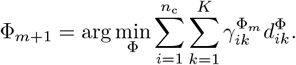

Substituting Eqs. (S10) and (S13) into the objective yields

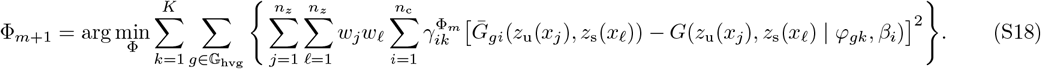

Because the objective in Eq. (S18) is separable across genes *g* and clusters *k*, the optimization decouples into independent subproblems. Specifically, each *φ*_*gk*_ can be updated by solving

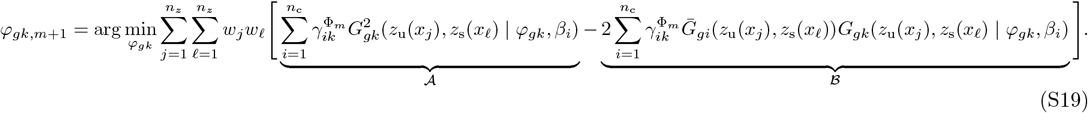

Both terms 𝒜 and ℬ involve summation over the entire cell population, implying that the computational cost scales linearly with *n*_c_. To reduce this cost, we approximate these sums using KDE.

Let

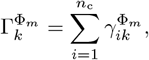

which is constant given Φ_*m*_. Define the normalized weights

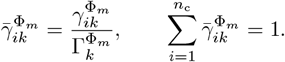

Since each 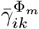 is associated with *β*_*i*_, these weights induce a probability distribution over *β*. Accordingly, the term 𝒜 can be approximated by

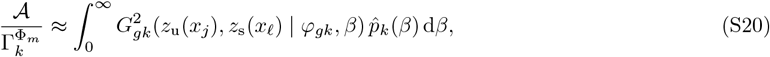

where the density 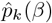 is estimated via KDE as

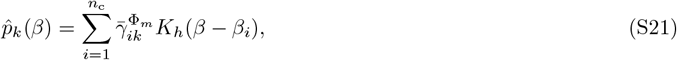

with kernel function *K*_*h*_ and bandwidth *h*.

The integral in Eq. (S20) is evaluated numerically using Gaussian-Legendre quadrature, yielding

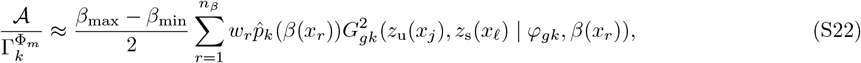

where

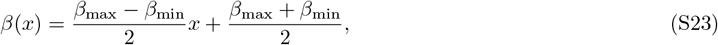

and 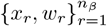 are quadrature nodes and weights computed using the `gausslegendre` function in Julia.

For the term ℬ, both 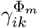 and 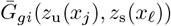 depend on *β*_*i*_. We therefore use the KDE trick and define a *z*-dependent density

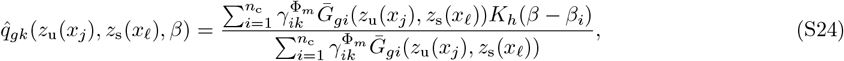

which satisfies normalization by construction.

Using this density, ℬ is approximated as

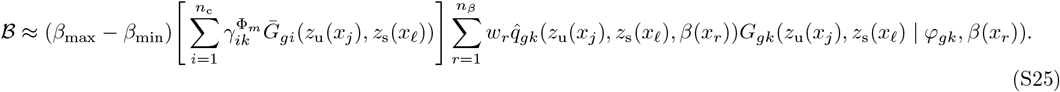

Combining Eqs. (S21)-(S25) allows efficient evaluation of the objective in Eq. (S19), with a computational cost that is independent of the number of cells *n*_c_. Parameter estimation for *φ*_*gk*_ thus reduces to the minimization of Eq. (S19), a step analogous to the maximization step of the classical EM algorithm. Each (*g, k*) subproblem is solved in parallel using the Nelder-Mead method implemented in the Julia package `Optim.jl`, selected for its robustness to initialization. The PGF-based majorization–minimization clustering procedure is summarized in Algorithm 1.

#### Algorithm 1

PRIME: PGF-based majorization-minimization clustering method.

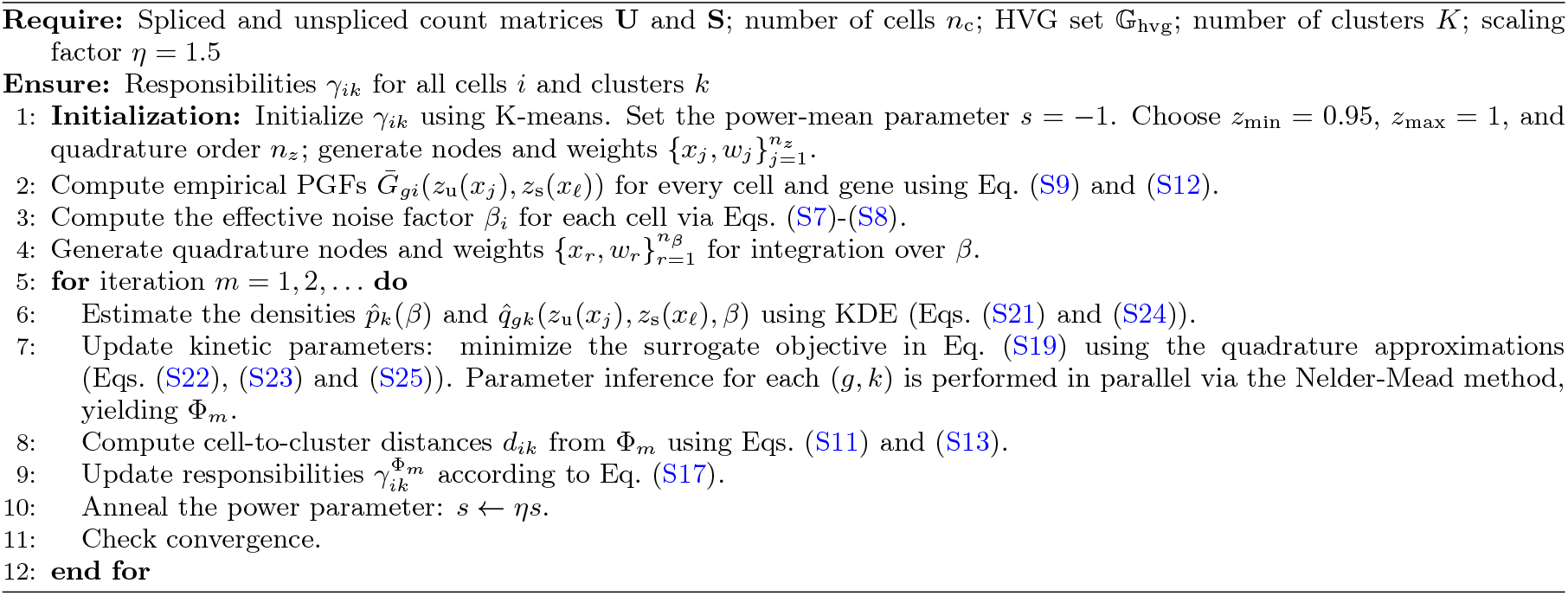

Given the inferred responsibilities, we next estimate the kinetic parameters for each gene within each cluster using the following algorithm.

#### Algorithm 2

PRIME: PGF-based Bayesian parameter inference method.

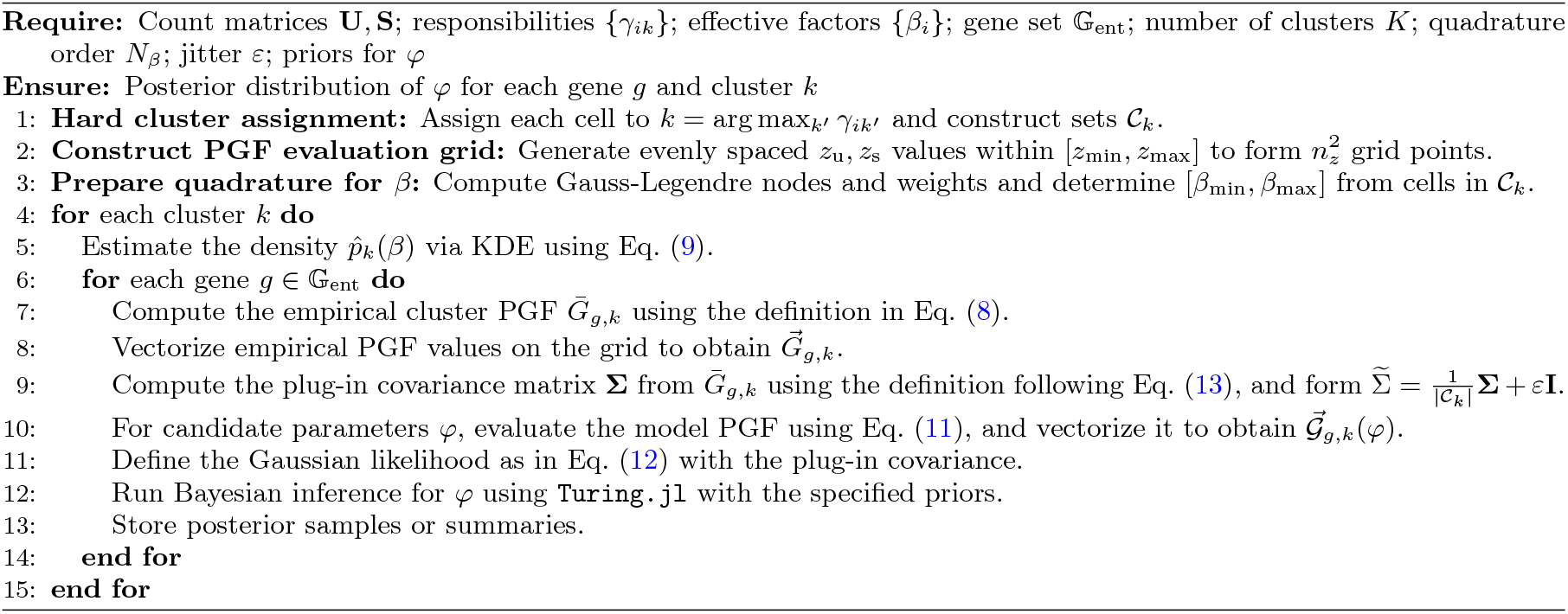

### Supplementary Note 4

#### Theorem (Asymptotic Normality of EPGF)

Let (*U*_1_, *S*_1_), (*U*_2_, *S*_2_), …, (*U*_*n*_, *S*_*n*_) be i.i.d. pairs of non-negative integer-valued random variables with a joint PGF 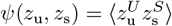. The bivariate EPGF is

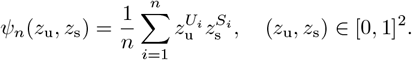

As *n* → ∞, the bivariate process 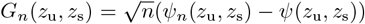 converges weakly to a zero-mean Gaussian process *G*(*z*_u_, *z*_s_) with covariance function:

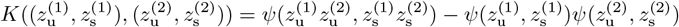

where 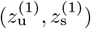 and 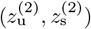 denote two pairs of evaluation values in PGF space.

*Proof*. The proof follows from the multivariate functional central limit theorem.

1. *Finite-Dimensional Convergence* For any finite collection of points 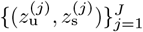 in the unit square, define the random vectors **Z** ∈ ℝ^*k*^ such that each component is 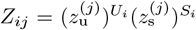. These vectors are i.i.d. with mean vector:

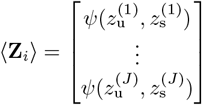

By the multivariate central limit theorem, the sum 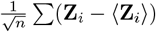 converges to a multivariate normal distribution *N* (**0**, Σ). The covariance between two indices *j* and *m* is:

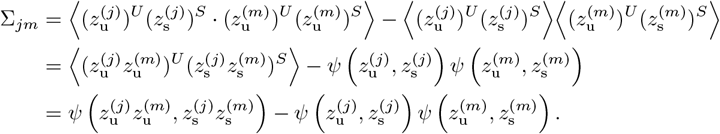
2. *Tightness* The class of bivariate functions 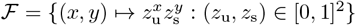 is a Donsker class. Since the partial derivatives with respect to *z*_u_ and *z*_s_ are bounded by 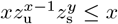 and 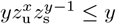 respectively, the envelope function *F* (*x, y*) = *x* + *y* is square-integrable if ⟨*U* ^2^ ⟩ *<* ∞ and ⟨*S*^2^ ⟩ *<* ∞. Thus, the sequence of processes *G*_*n*_(*z*_u_, *z*_s_) is tight in *C*([0, 1]^2^). By Prohorov’s Theorem [89], the finite-dimensional convergence and tightness imply weak convergence to the Gaussian process *G*(*z*_u_, *z*_s_).

**FIG. S1.**
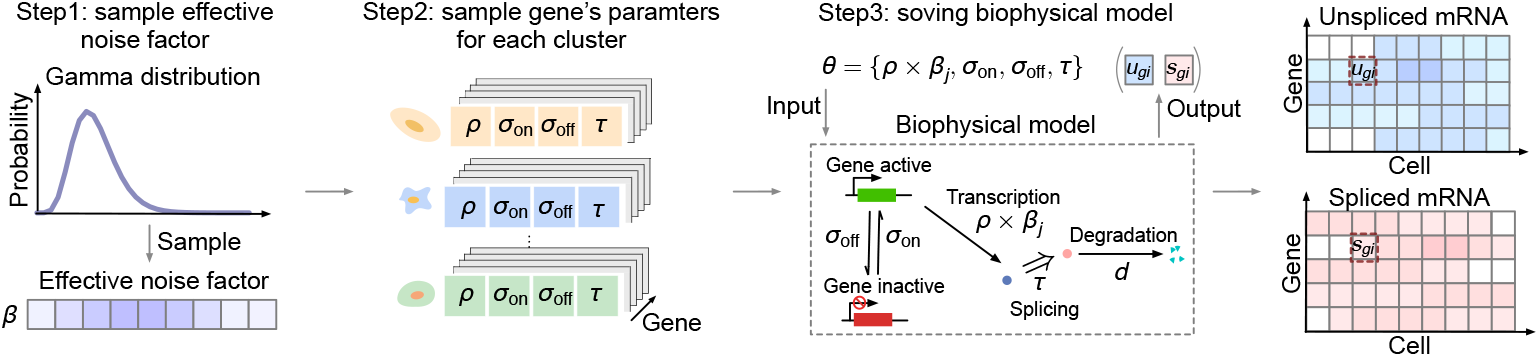
Workflow for synthetic data generation. Each cell is assigned an effective noise factor *β* sampled from a Gamma distribution to model extrinsic and technical noise. Cluster structure and gene-specific kinetic parameters are specified, and unspliced and spliced RNA counts in steady-state conditions are generated by stochastic simulation of the transcription model (Eq. (1)).

**FIG. S2.**
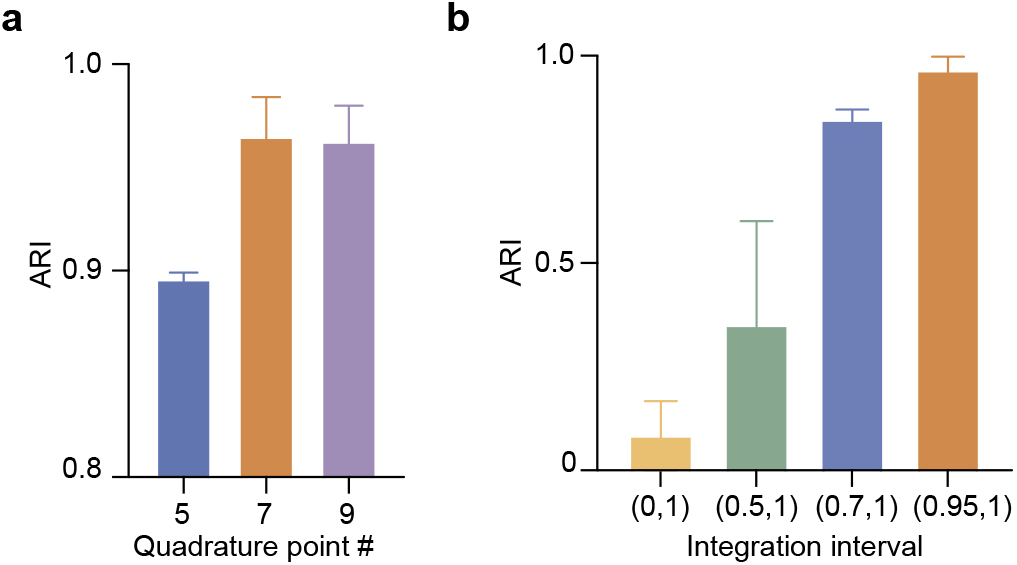
Sensitivity analysis of quadrature resolution and integration domain. (a) Clustering accuracy (ARI) as a function of the number of Gauss–Legendre quadrature points *n*_*z*_ ∈ {5, 7, 9} evaluated on a synthetic dataset with ten clusters. Performance stabilizes at *n*_*z*_ = 7, indicating sufficient resolution without additional computational cost. (b) Clustering accuracy under different integration domains in PGF space: (0, 1), (0.5, 1), (0.7, 1), and (0.95, 1). Restricting the domain to (0.95, 1) consistently yields the highest accuracy. Unless otherwise stated, PRIME uses *n*_*z*_ = 7 and the integration range (0.95, 1). Bars represent the mean, and error bars indicate the SEM across three independent runs.

**FIG. S3.**
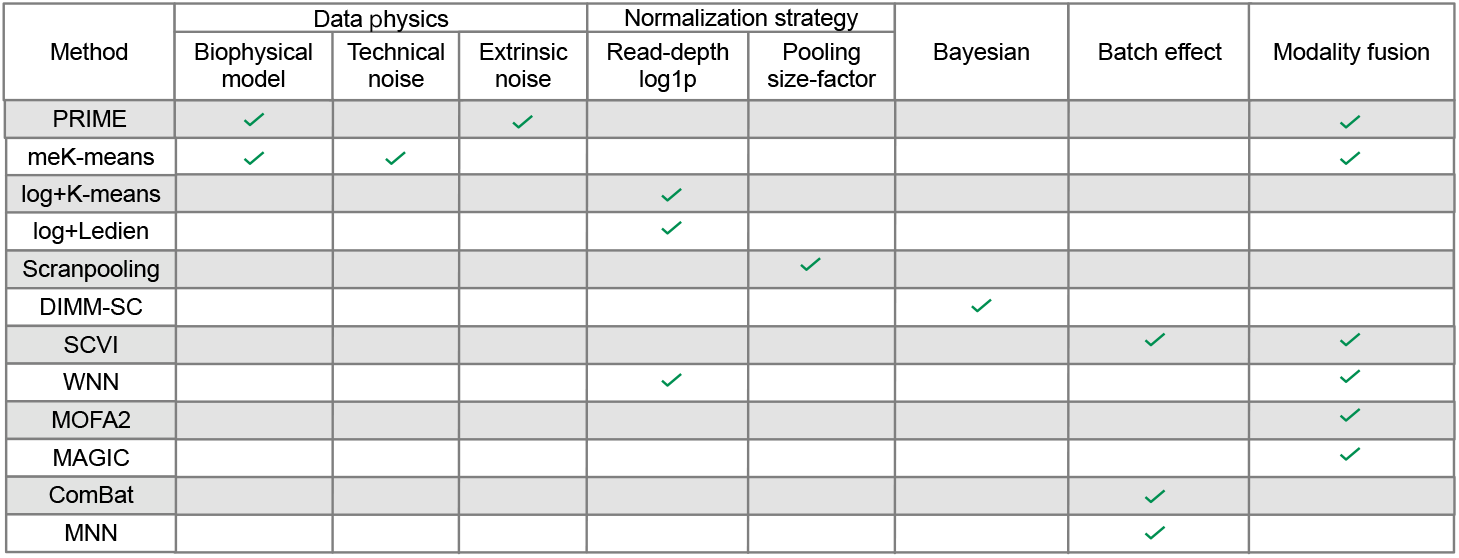
Functional comparison of clustering methods included in the benchmarking analysis. Summary of the methodological features of all clustering approaches evaluated in this study. Methods are grouped according to their primary design principles, including biophysical model-driven inference, normalization or noise-correction strategies, batch-effect correction, multimodal integration, and Bayesian or probabilistic frameworks. The table highlights which functional components are present in each method, providing a unified overview of the methodological landscape used for comparison with PRIME.

**FIG. S4.**
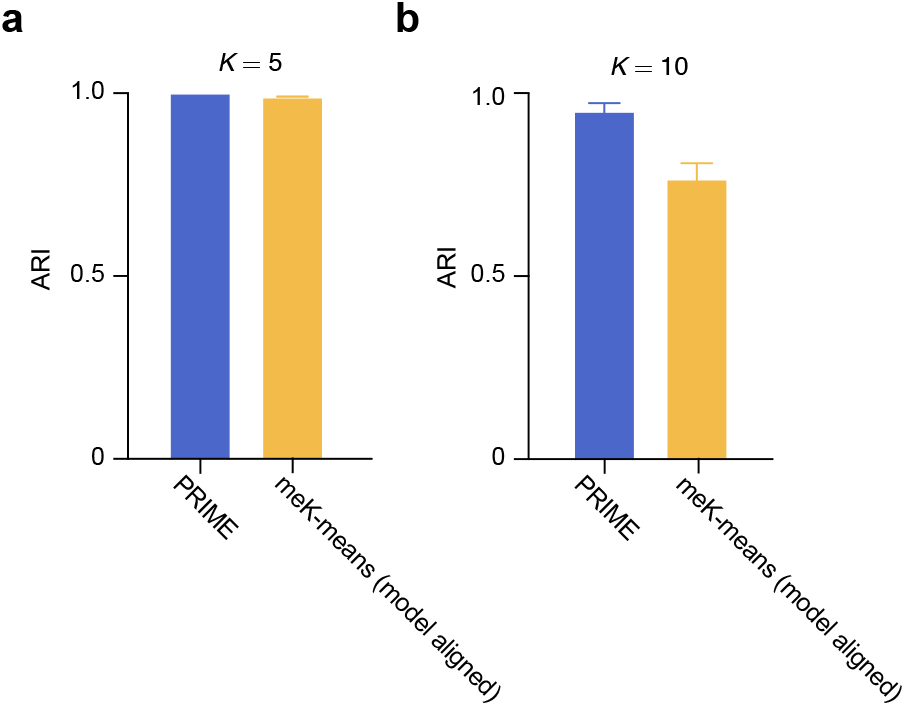
Clustering performance, measured by ARI, comparing PRIME and meK-means when the meK-means model is aligned with the data-generating model used to simulate the synthetic data. (a) Results for the synthetic dataset with *K* = 5 clusters, where PRIME and meK-means achieve comparable clustering accuracy. (b) Results for the dataset with *K* = 10 clusters, where meK-means underperforms PRIME. Bars represent the mean, and error bars indicate the SEM across three independent runs.

**FIG. S5.**
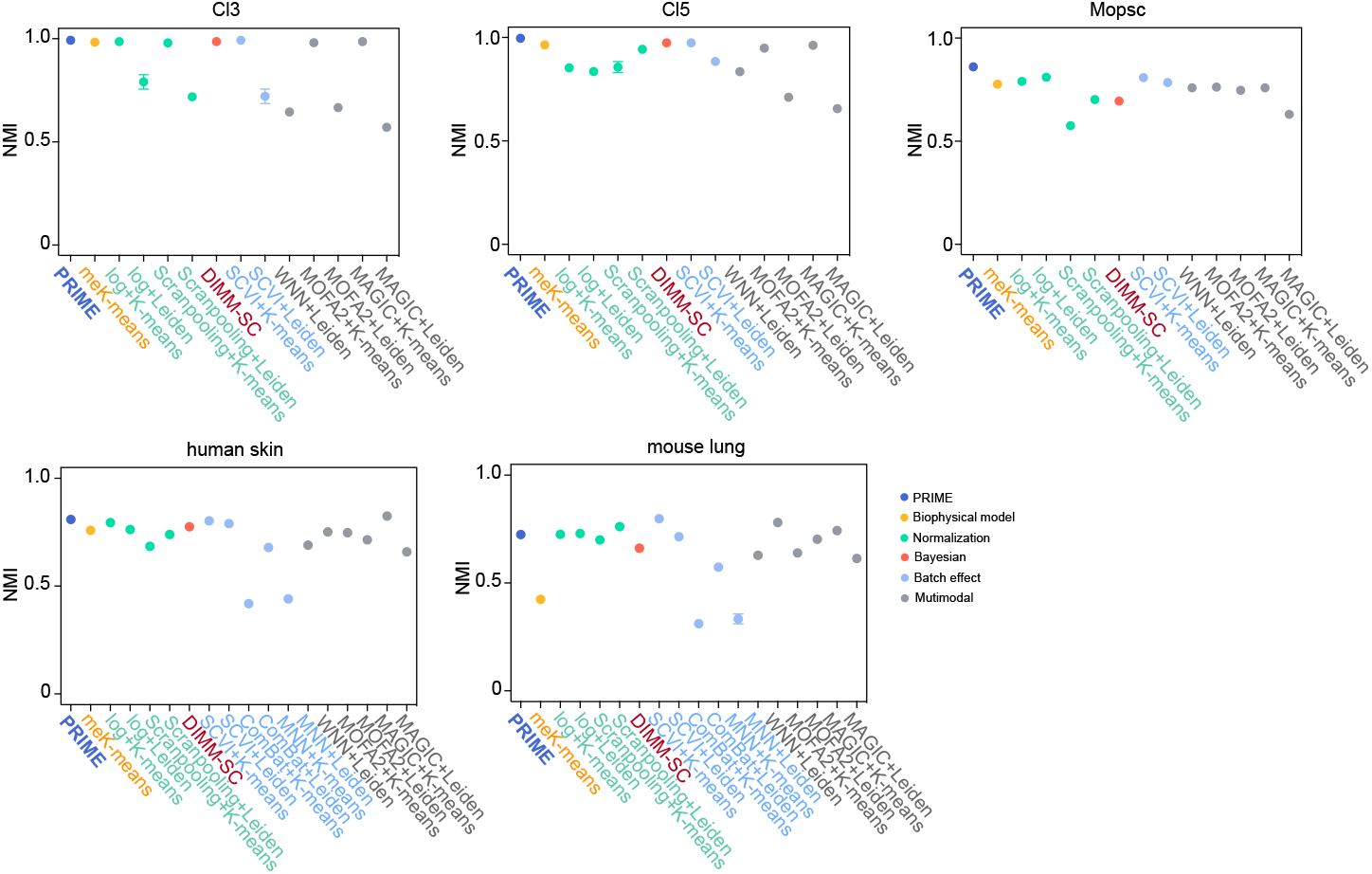
Additional benchmarking of PRIME on experimental datasets. Clustering performance of PRIME and competing methods evaluated across five experimental datasets. Accuracy is quantified using Normalized Mutual Information (NMI). Methods include normalization-based (log transformation, Scran pooling), batch-corrected (ComBat, MNN, SCVI), multimodal-integration (MAGIC, MOFA2, WNN), Bayesian (DIMM-SC), and mechanistic (meK-means) approaches. Dots represent the mean across five independent runs, and error bars denote the SEM. PRIME consistently ranks among the top-performing methods across datasets.

**FIG. S6.**
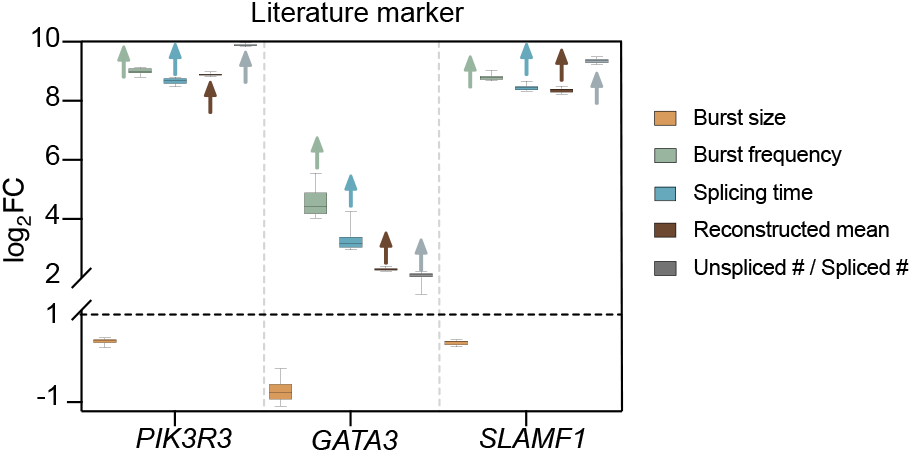
Kinetic signatures of CTCL-associated genes identified by PRIME. Reconstructed expression and inferred kinetic parameters (burst size, burst frequency, and splicing time) for *PIK3R3, GATA3*, and *SLAMF1* in `CTCL` and `ctrl` samples. All three genes display the characteristic kinetic pattern identified in Fig. 6f, supporting their association with CTCL.

**TABLE S1.** Ground-truth kinetic parameter values for the first 10 genes in a cluster of 1000 cells from the *K* = 5 synthetic dataset.

| Gene index | $\rho$ | $\sigma_{\text{on}}$ | $\sigma_{\text{off}}$ | $\tau$ |
| --- | --- | --- | --- | --- |
| 1 | 7.24834 | 0.910108 | 1.68449 | 0.821993 |
| 2 | 4.91289 | 1.01182 | 2.18204 | 1.99910 |
| 3 | 3.80678 | 1.06367 | 1.44434 | 1.68821 |
| 4 | 5.38383 | 0.908455 | 1.86954 | 1.46530 |
| 5 | 5.08997 | 1.16958 | 1.13977 | 1.23440 |
| 6 | 4.24687 | 0.887943 | 1.10879 | 1.61312 |
| 7 | 2.75499 | 0.960232 | 1.71333 | 0.551839 |
| 8 | 4.90084 | 0.977727 | 1.84107 | 0.771582 |
| 9 | 25.4514 | 1.04356 | 1.34934 | 1.26972 |
| 10 | 4.10571 | 1.18987 | 3.44665 | 1.28086 |

